# Tau pathology spreads between anatomically-connected regions of the brain and is modulated by a LRRK2 mutation

**DOI:** 10.1101/2020.10.13.337857

**Authors:** Michael X. Henderson, Eli J. Cornblath, Howard L. Li, Lakshmi Changolkar, Bin Zhang, Hannah J. Brown, Ronald J. Gathagan, Modupe F. Olufemi, Danielle S. Bassett, John Q. Trojanowski, Virginia M.Y. Lee

## Abstract

Tau pathology is a diagnostic feature of Alzheimer’s disease (AD) but is also a prominent feature of Parkinson’s disease (PD), including genetic forms of PD with mutations in leucine-rich repeat kinase 2 (*LRRK2*). In both diseases, tau pathology is progressive and correlates with cognitive decline. Neuropathological staging studies in humans and mouse models have suggested that tau spreads through the brain, but it is unclear how neuroanatomical connections, spatial proximity, and regional vulnerability contribute to pathology spread. Further, it is unknown how mutations in the *LRRK2* gene may modulate susceptibility to tau pathology’s initiation or spread. In this study, we used seed-based models of tauopathy to capture spatiotemporal patterns of pathology in mice. Following the injection of AD brain-derived tau into the brains of non-transgenic mice, tau pathology spreads progressively through the brain in a spatiotemporal pattern that is well-explained by anatomical connectivity. We validated and compared network models based on diffusion along anatomical connections to predict tau spread, estimate regional vulnerability to tau pathology, and investigate gene expression patterns related to regional vulnerability. We further investigated tau pathology spread in mice harboring a mutation in LRRK2 and found that while tau pathology spread is still constrained by anatomical connectivity, it spreads preferentially in a retrograde direction to regions that are otherwise resilient in wildtype mice. This study provides a quantitative demonstration that tau pathology spreads along anatomical connections, explores the kinetics of this spread, and provides a platform for investigating the effect of genetic risk factors and treatments on the progression of tauopathies.

## INTRODUCTION

Neurodegenerative diseases, including Parkinson’s disease (PD) and Alzheimer’s disease (AD), are estimated to affect over 60 million people worldwide ^1,2^. Neurological symptoms and the presence of pathological protein inclusions are used to categorize the two diseases. However, there exists substantial overlap in both symptoms and pathologies, especially as these diseases progress ^3,4^ Tau pathology, while diagnostic of AD and other primary tauopathies, appears prominently in PD, PD dementia (PDD), and dementia with Lewy bodies (DLB), where it correlates with α-synuclein pathological burden and cognitive decline ^5,6^ These data suggest that multiple pathologies may act additively to influence disease progression, and that underlying risk factors for one pathology may confer risk for additional pathologies.

Multiple factors, including lifestyle, exposure to environmental pathogens, and genetic variants can influence risk of developing neurodegenerative disease. Up to 27% of PD has been estimated to be heritable ^7,8^ through both rare mutations and more common polymorphisms in the genome. The most common cause of familial PD and a common risk factor for idiopathic PD is a mutation in the gene encoding leucine-rich repeat kinase 2 (LRRK2) ^9^. The most prevalent of these mutations, p.G2019S, confers a 25-42.5% risk of developing PD ^10^. Although these patients show similar symptoms to idiopathic PD, neuropathologically, 21-54% of patients lack the hallmark α-synuclein Lewy bodies exhibited by idiopathic PD patients ^11–13^. Notably, the majority of *LRRK2* mutation carriers exhibit tau pathology ^13,14^ The tau pathology in idiopathic PD and *LRRK2*-PD is similar in conformation and distribution to AD tau and may be partially responsible for the cognitive decline seen in these patients during the disease course ^6,13^. The appearance of tau pathology in *LRRK2* mutation carriers suggests that the genetic risk conferred by *LRRK2* mutations may alter the development or progression of tau pathology. If *LRRK2* mutations do in fact modulate tau pathology, then LRRK2 kinase inhibitors being developed for the treatment of genetic and idiopathic PD may influence tau pathology in these patients, and possibly in AD and other tauopathy patients as well.

In the current study, we investigated processes underlying the development and spread of tau pathology, including the impact of the p.G2019S *LRRK2* mutation and LRRK2 kinase inhibitors. Our initial experiments performed in primary neurons from non-transgenic (NTG) and LRRK2^G2019S^ mutation-bearing neurons revealed that there is minimal impact of the LRRK2 mutation or LRRK2 kinase inhibition on the acute development of tau pathology. Therefore, we designed and implemented a series of experiments aimed first at developing a deeper understanding of how tau pathology spreads through the mouse brain following an intracranial injection of pathological tau, and second how LRRK2^G2019S^ modulates the spread of tau pathology. Consistent with hypotheses from human neuropathology studies, we found that tau spreads through the mouse brain in a constrained spatiotemporal pattern that can be predicted from neuroanatomical connectivity. We further demonstrate that anterograde and retrograde connectivity both contribute to pathology spread and a model incorporating both directions shows the highest predictivity. Understanding the nature of spread along anatomical connections also enabled us to estimate regional vulnerability to tau pathology, and to perform an unbiased assessment for genes that may underlie this regional vulnerability. Finally, we performed similar quantitative pathology and computational modeling to understand differences in tau pathology spread between LRRK2^G2019S^ and NTG mice. LRRK2^G2019S^ mice exhibit a bias towards retrograde spread of tau pathology, providing insight into the network-level impact of cell biology events. This work provides a framework for understanding the spread of pathological tau throughout the brain, investigating the impact of genetic risk factors, and assessing the impact of therapeutic interventions related to LRRK2.

## RESULTS

### The G2019S LRRK2 mutation does not exacerbate tau pathology in primary neurons

Primary neuron cultures provide a rapid and accessible method for the evaluation of cellular phenotypes and the efficacy of therapeutics. We have recently developed primary neuron models which develop insoluble aggregates of tau pathology without overexpression of tau to allow the assessment of compounds which may modify tau pathology 15. For the current study, we were particularly interested in evaluating whether or not the LRRK2^G2019S^ mutation and LRRK2 kinase inhibition would modify tau pathology in primary neurons. We first evaluated the ability of three LRRK2 inhibitors—PF-06447475 (PF-475), PF-06685360 (PF-360), and MLi-2 to inhibit LRRK2 kinase activity in LRRK2^G2019S^ primary hippocampal neurons that overexpress LRRK2^G2019S^ under the endogenous mouse promoter ^16^. Total LRRK2 and pS935 LRRK2 were monitored as a common and reliable readout for LRRK2 kinase activity ^17–20^. All three compounds were able to reduce pS935 LRRK2 levels strongly at low nanomolar concentrations with minimal disruption of total LRRK2 levels (Fig. 1A–1C).

**Figure 1.**
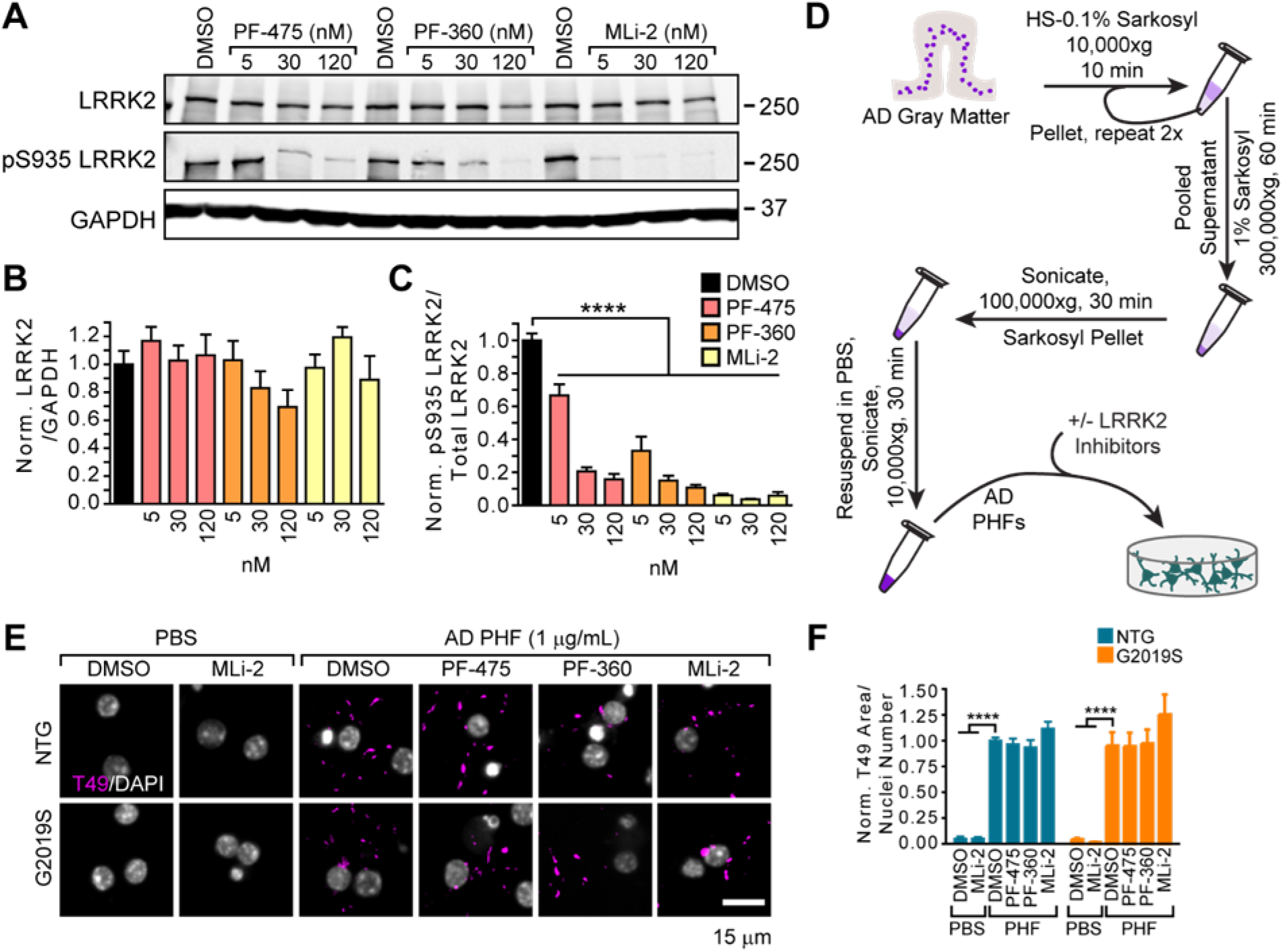
The G2019S *Lrrk2* mutation does not exacerbate tau pathology in primary neurons. **(A)** Primary cortical neurons from LRRK2^G2019S^ mice were treated at 5 days *in vino* (DIV) with LRRK2 inhibitors at the noted concentrations and lysate was harvested at 21 DIV. Western blot is shown for total LRRK2 and pS935 LRRK2. the latter being a proxy for LRRK2 activity. (B) Quantification of total LRRK2 normalized to GAPDH and vehicle treatment. (C) Quantification of pS935 normalized to total LRRK2 and vehicle treatment. (D) Schematic diagram of AD PHF tau extraction from human brain tissue and subsequent treatment of primary neurons. (E) Primary hippocampal neurons from NTG or LRRK2^G2019S^ mice were treated with vehicle or LRRK2 inhibitors at 300 hM at 5 DIV. They were further treated with AD PHF tau at 1 μg/mL at 7 DIV. and both fixed and stained for insoluble mouse tau (T49. magenta) and DAPI (gray) at 21 DIV. Scale bar =15 μm. (F) Quantification of the insoluble tau area/nuclei number in each condition. No insoluble tau was present in the absence of PHF treatment and there was no overall effect of genotype (Two-way ANOVA; genotype effect p=O.837. Dunnett’s multiple comparison test within genotype: NTG: DMSO-PHF vs. DMSO ****p<0.0001; DMSO-PHF vs. MLi-2 ****p<0.0001; G2019S: : DMSO-PHF vs. DMSO ****p<0.0001; DMSO-PHF vs. MLi-2 ****p<0.0001; all other values were not statistically significant). Data are represented as mean ± SEM.

We then utilized a biochemical sequential detergent extraction of gray matter from AD patient brains to obtain an enriched fraction of paired helical filament (PHF) tau (Fig. 1D) 15. This extraction method yielded a final purity of 16.1-35.7% tau, with 0.01% or less α-synuclein and amyloid β (Fig. S1A). This purified tau fraction retains the pathogenic conformation present in human disease and induces the misfolding of tau in primary neurons and mice without the overexpression of tau 15; this AD tau therefore serves as a valuable tool to assess tau pathology in different genetic backgrounds. Primary hippocampal neurons from NTG or LRRK2^G2019S^ mice were treated with vehicle or LRRK2 inhibitors followed by treatment with DPBS or PHF tau (Fig. 1E, 1F). Notably, neither NTG nor LRRK2^G2019S^ neurons form tau inclusions without the addition of PHF tau. However, 14 days post-treatment with PHF tau, neurons form detergent-insoluble tau inclusions from endogenous mouse tau, which do not differ by neuron genotype. Further, LRRK2 kinase inhibition had no effect on tau pathology in these neurons (Fig. 1E, 1F).

We also evaluated the effect of LRRK2^G2019S^ and LRRK2 kinase inhibition in a second primary neuron model of tauopathy (Fig. S1B). In this model, recombinant tau is shaken, sonicated and passaged into monomer sequentially in a manner that stochastically generates fibrillar tau without the need for a co-factor such as heparin 15. These “X-T40” pre-formed fibrils (PFFs) can be sonicated and added to NTG primary neurons and will seed the misfolding of endogenous tau. Neither NTG, nor LRRK2^G2019S^ neurons have detectable levels of pS202/T205 tau without the addition of X-T40 PFFs, but PFFs induce robust tau inclusions in neurons (Fig. S1B, S1C). Interestingly, we observed a slight reduction of tau pathology in LRRK2^G2019S^ neurons, with no apparent effect of LRRK2 kinase inhibition (Fig. S1B). There was a small reduction in overall dendritic area (MAP2) with PFF treatment that was reversed with the highest dose of PF-360 (Fig. S1D). Together, these results suggest that LRRK2^G2019S^ and LRRK2 kinase inhibition have a minimal effect on the cell-autonomous production of tau pathology in neurons. However, primary neuron cultures do not allow for the assessment of the impact of LRRK2 on tau pathology over long time periods, effects on cell-to-cell spread, or non-cell autonomous effects.

### Quantitative immunohistochemistry to evaluate pathological tau spread

Although previous work had been done crossing LRRK2 transgenic mice to transgenic mice that overexpress tau with mutations (Mikhail et al., 2015; Nguyen et al., 2018), these studies are not equipped to assess tau spread. To assess this possibility, we utilized a novel seed-based model of tauopathy in which injection of PHF tau into NTG mice can induce the misfolding of endogenous tau into hyper-phosphorylated tau inclusions in a time and region-dependent manner 15. Previous studies have suggested that this spatiotemporal pattern of tau pathology induction is consistent with spread along neuroanatomical connections ^15,21^, although these patterns were not quantitatively evaluated. To understand how tau pathology spreads through the brain and how spread is modified by the G2019S mutation in LRRK2, NTG and LRRK2^G2019S^ mice were injected with PHF tau in the hippocampus and overlaying cortex (Fig. 2A). Mice were allowed to age 1, 3, 6, or 9 months post-injection (MPI) to capture the temporal dynamics of tau pathology spread (Fig. 2B). Brains were then sectioned, and representative sections (Fig. 2C) were selected and stained for phosphorylated tau pathology throughout the brain. We manually annotated 194 regions on the selected sections (Fig. 2D, Fig. S2) so that pathology could be quantified as the percentage of each area occupied by pathology (Fig. 2E, 2F). To mitigate overinterpretation of individual sections, a second set of nearby sections were similarly annotated and quantified (16,684 annotations total), and the average for each region was used for subsequent analyses.

**Figure 2.**
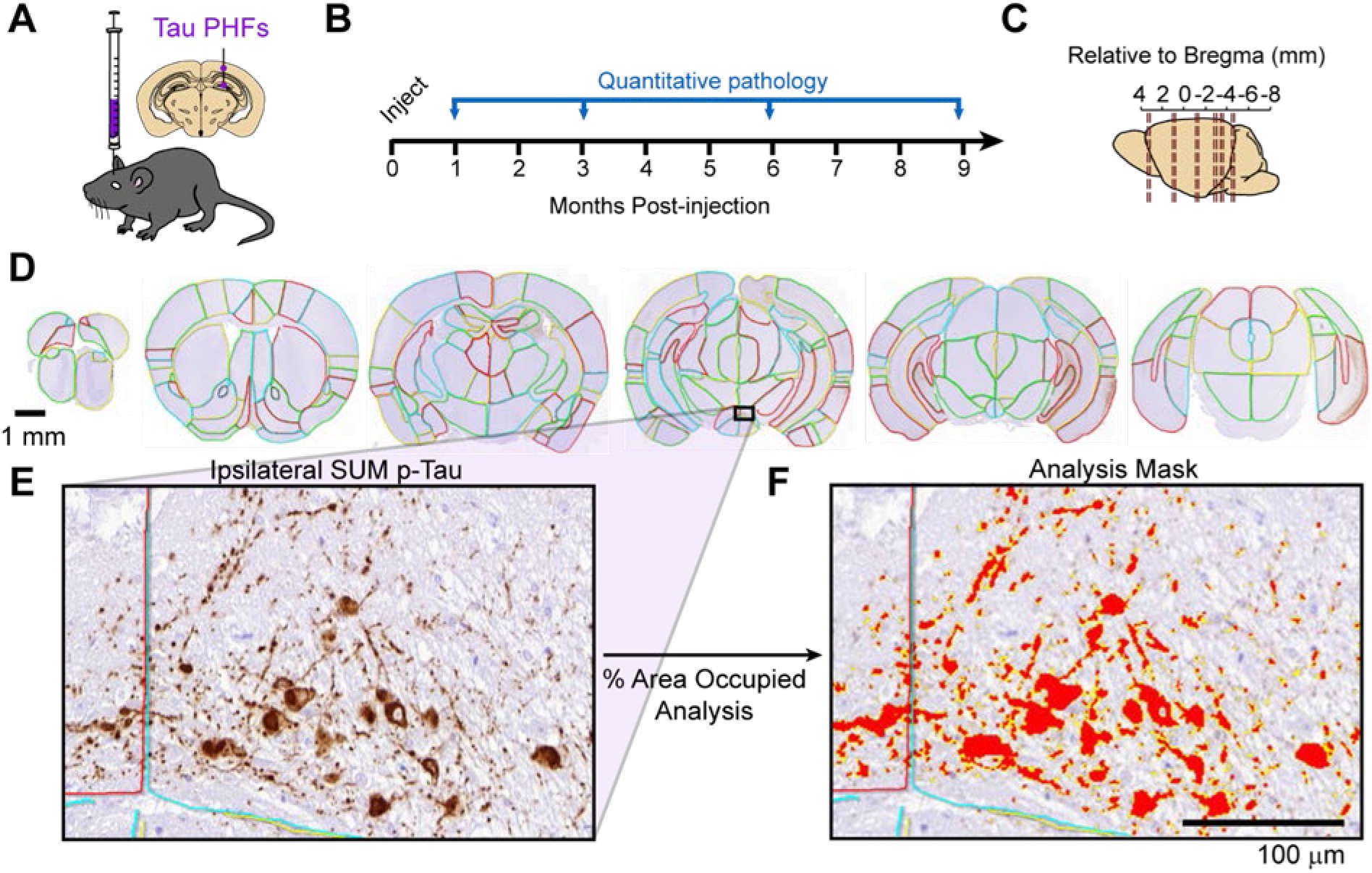
Quantitative immunohistochemistry to evaluate pathological tau spread. **(A)** Non-transgenic mice were injected unilaterally with AD PHF tau in the hippocampus and overlaying cortex as shown. (B) Mice were sacrificed 1, 3, 6, or 9 months following injection. (C) Mouse brain were sectioned, and the sections representing the regions shown were stained for pathological tau. (D) Representative sections were selected and 194 regions were annotated for each brain. A second set of nearby sections was similarly annotated to reduce selection bias. Scale bar = 1 mm. Annotation colors are arbitrary. (E) An enlarged image of the annotated supramammillary nucleus (SUM) is shown with the inclusions stained for pS2O2T2O5 tau. (F) Annotations allow automated quantification of % area occupied with pathology in specific regions of the brain. An analysis mask is overlaid on the image in (E) to demonstrate this quantification of pathology. Scale bar =100 μm.

### Quantitative pathology mapping reveals dynamic patterns of tau pathology spread over time

The use of quantitative pathology allows an assessment of patterns of tau pathology across the brain over time and within individual regions. Tau pathology for each region was averaged across mice at each time point, providing pseudo-longitudinal measures of tau pathology. These measures were plotted on anatomical heat maps of the mouse brain (Fig. 3A) and as a region-by-time matrix (Fig. S3). Several whole-brain patterns are apparent. At 1 MPI, there is minimal tau pathology outside of the injection sites. At 3 MPI, more pathology accumulates ipsilateral to the injection site including hippocampal and entorhinal regions. By 6 MPI, pathology has spread to the contralateral hippocampus and associated cortical areas. By 9 MPI, pathology has continued to spread, affecting more rostral regions, although certain rostral cortical and contralateral thalamic regions show minimal pathology. While these global patterns are indicative of tau pathology spread, it is also informative to consider temporal patterns in individual regions (Fig. 3B). For example, some regions, like the supramammillary nucleus (iSUM) show a linear increase in pathology over time. Other regions, like the contralateral CA1 region of the hippocampus (cCA1), have minimal pathology until 6 MPI. The accessory olfactory bulb (iAOB) has minimal pathology until 9 MPI. In contrast, the iDG at the site of injection has rapid induction of pathology, which stabilizes after 3 MPI. Representative images of all regions are available in the supplement (Fig. S9). Together these patterns are highly suggestive of tau pathology spread throughout the brain, which may involve transmission along neuroanatomical connections or other regional factors.

**Figure 3.**
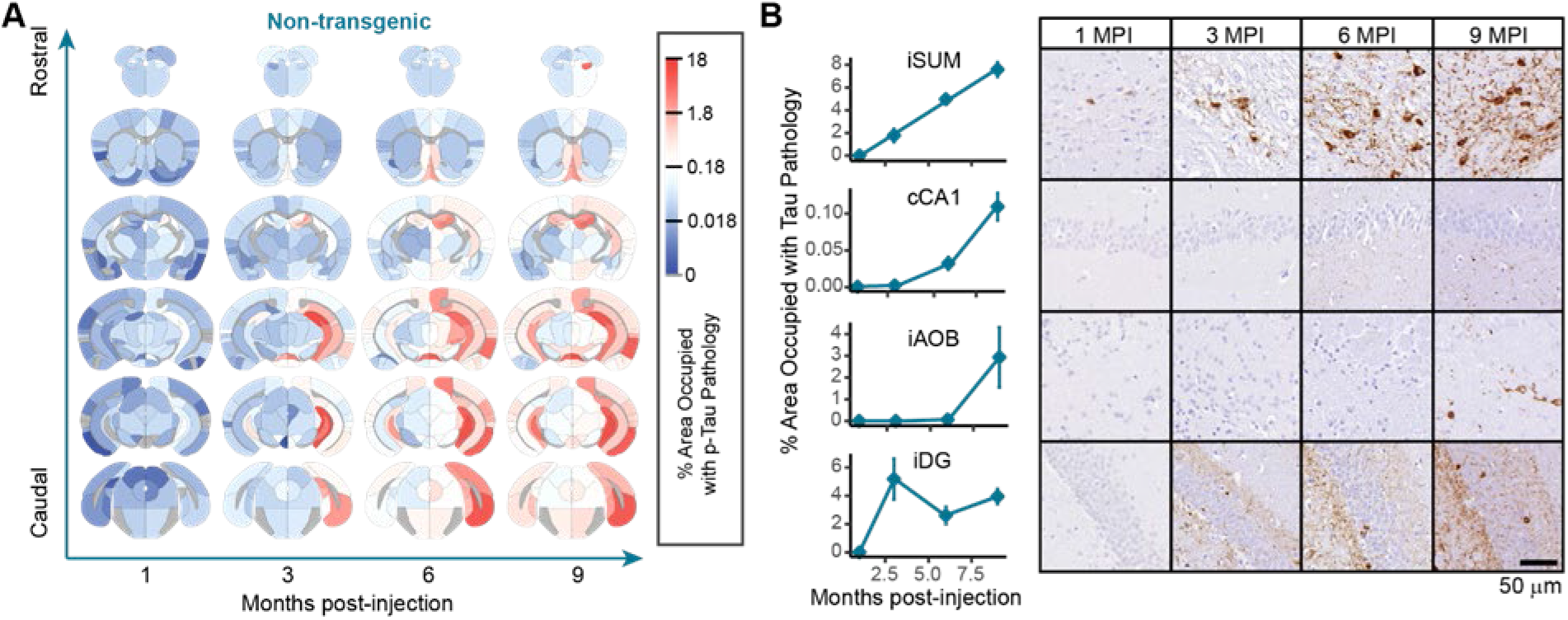
Quantitative pathology mapping reveals dynamic patterns of tau pathology spread over time. (A) Regional pathology measures plotted on anatomical maps as a heat map. with blue representing minimal pathology, white representing moderate pathology, and red representing substantial pathology. Note that tau pathology was not quantified in white matter regions, and these regions are gray. (B) The % of area occupied with tau pathology is plotted as a function of time for 4 different regions (iSUM. cCAl. iAOB, iDG) demonstrating four distinct pattern of pathology propagation. Many of the other regions also fell into one of these patterns. The iSUM shows an almost linear increase in pathology over 9 months, beginning at 1 MPI. The cCAl has only minimal pathology up to 3 months, after which neuritic pathology begins to accumulate at 6 and 9 MPI. The iAOB has minimal pathology until 9 MPI. showing an extensive delay following injection. Finally, the iDG (an injection site) rapidly accumulates pathology at 3 MPI, but pathology is not further elevated at 6 or 9 MPI. Representative images of the regions plotted in the left panel are stained for p-tau and directly adjacent to the plots demonstrating the pathology patterns. Representative images for all regions can be found in the Supplementary Material. Scale bar = 50 μm.

### Tau pathology spread is predicted by diffusion through the neuroanatomical connectome

There is substantial evidence that tau can be released from neurons ^22–24^ in an activitydependent manner ^25,26^ and internalized by other neurons ^27–30^ likely leading to the spread of tau pathology throughout the brain. However, it is still unclear precisely how spread occurs. The main hypotheses are that tau pathology spreads between anatomically connected regions of the brain, between physically contiguous regions, or to selectively vulnerable populations of neurons. Of course, these mechanisms of spread are not mutually exclusive; thus, we sought to assess how each of these putative routes contribute to the observed pathology spread in mice.

We evaluated the ability of a network diffusion model with neuroanatomical connectivity (brain-map.org; ^31^ as a scaffold to predict the empirical measures of tau pathology over time. Allen Brain Atlas (ABA) regions at the injection sites (iDG, iCA1, iCA3, iRSPagl, iVISam) were used as seed regions for initiation of the model. Direct connectivity of these regions was highest in hippocampal, septal and entorhinal regions of the brain (Fig. S4A), and direct connectivity to the injection site was highly correlated with tau pathology measures in those regions (Fig. S4B, S4C). Our model posits that tau spreads from the injection site along anatomical connections both in retrograde and anterograde directions, with the final amount of regional pathology determined by a weighted sum of these two independent processes. This bidirectional anatomical connectivity model weakly predicted tau pathology at 1 MPI, likely due to minimal spread at this time point, and strongly predicted tau pathology at 3, 6, and 9 MPI (Fig. 4A, 4D). Additionally, this bidirectional anatomical connectivity model outperformed a model in which tau spread was proportional to the Euclidean distance between regions (Fig. 4B, 4D). In the Euclidean distance plots, the 5 outlying regions in the upper right were the injection sites, as noted (Fig. 4B); however, inclusion or exclusion of these sites during model optimization did not measurably affect the fit of the Euclidean distance model (Fig. S5). Rewiring the network to preserve in-degree (Fig. S6A) or out-degree connectivity (Fig. S6B) also reduced model performance. To further validate the anatomical connectivity model, we ensured that the model’s performance was specific to the choice of the experimental injection site, compared with 500 randomly chosen sets of 5 regions with mean spatial proximity similar to that of the experimental injection sites (Fig. 4C). Models using the experimental seed regions were among the best performing models at all time points (Fig. 4C), confirming the specificity of our model to the experimental injection site. Interestingly, the performance of models using random sites could be partially explained by their topological and geometric similarity with the experimental sites (Fig. S6C).

**Figure 4.**
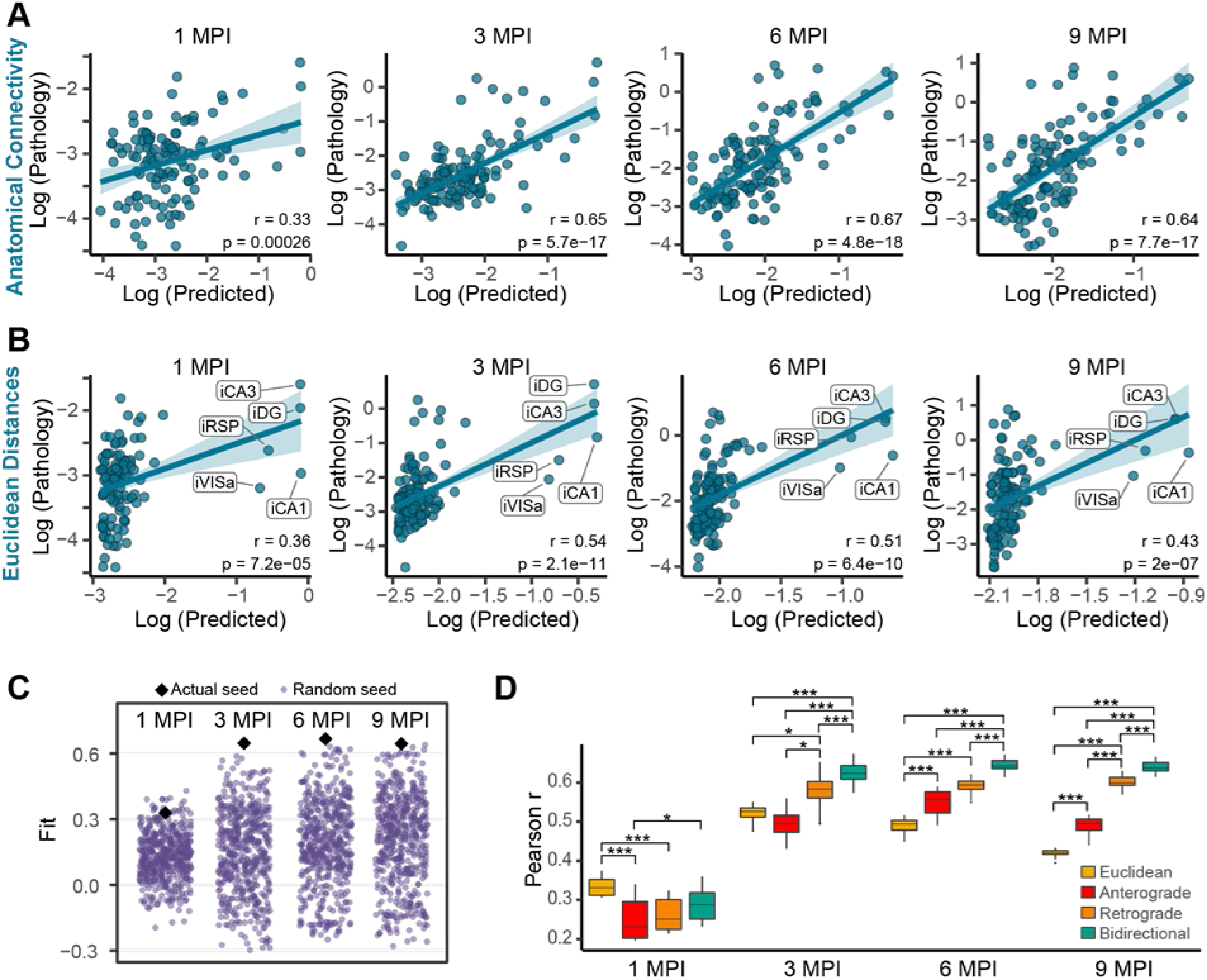
Tau pathology spread is predicted by diffusion through ueuroauatomical connectivity. (A-B) Predictions of log tau pathology (*x*-axis) from spread models based on (A) retrograde and anterograde anatomical connections or (B) Euclidean distance, plotted against log actual regional tau pathology values (*y*-axis) at 1. 3. 6, and 9 MPI. The solid lines represent the lines of best fit and the shaded ribbons represent the 95% prediction intervals. The r and p values for the Pearson correlation between model fitted values and observed pathology are noted on the plots. In (B). the injection site regions are noted, but did not impact model fit (Fig. S5). (C) To evaluate the specificity of the seed sites to predict the pathology spread pattan. 500 alternate combinations of 5 seed sites (purple dots) were evaluated for their ability to predict pathology spread at 1, 3, 6. and 9 MPI. Using the actual 5 injection regions (black diamonds) produced among the best fits at all time points (1 MPI: p=0.042, 3 MPI: p<0.002, 6 MPI: p<0.002, 9 MPI: p<0.002. (D) Distributions of model fits in 500 held-out samples using Euclidean, anterograde, retrograde, and bidirectional models. Fit differences were analyzed by pairwise two-tailed non-parametric tests for different models. All connectivity models outperform Euclidean distance after 3 MPI. and a bidirectional model outperforms either a retrograde or anterograde model alone (*p<0.05. **p<0.01. ***p<0.002).

Finally, we rigorously evaluated the out-of sample performance of the bidirectional anatomical connectivity model compared to three other models in which spread was based on either Euclidean distance, anterograde spread alone, or retrograde spread alone. In order to compare the distributions of out-of-sample fits between each of the 4 models, we generated 500 train-test splits of the mice used in the study. Next, we obtained model parameters (time constants and regression weights for anterograde and retrograde spread) in the training set, and evaluated model performance in the test set at each time point. Our measure of performance was the spatial Pearson correlation coefficient between the observed pathology in the test set and the predicted pathology estimated by each model. This analysis revealed that the bidirectional model was superior to both the anterograde and retrograde models, and that all 3 connectivity-based models were superior to the Euclidean distance model by 6 and 9 MPI (Fig. 4D). In summary, these findings provide strong support for the notion that both anterograde and retrograde spread along anatomical connections independently contribute to the propagation of tau pathology.

### Model residuals as an estimate of regional vulnerability

A third possible mediator of tau pathology spread is intrinsic neuronal vulnerability. While this concept is less easily parsed for tauopathies than for α-synucleinopathies where dopaminergic neurons appear particularly vulnerable, our network model of tau pathology spread can be used to infer regional vulnerability. We have previously validated this approach by showing that residuals from network model predictions of pathology correlated with α-synuclein gene expression, which was expected to contribute to regional vulnerability ^32^. Here, we estimated regional vulnerability to tau pathology spread by taking the residuals from the bidirectional anatomical connectivity model and averaging them over 3, 6, and 9 MPI (Fig. 5A). We excluded 1 MPI due to the low model predictivity and dissimilarity to residuals from other time points (Fig. S7A). This estimate of regional vulnerability revealed that amygdalar, thalamic, and rostral cortical nuclei were resilient, while septal, mesencephalic, and caudal cortical regions were more vulnerable (Fig. 5A). We then tested whether tau gene (*Mapt*) expression from ABA *in situ* hybridization data was associated with vulnerability. *Mapt* is expressed quite broadly in the brain, including most gray matter and white matter regions, and showed no association with regional vulnerability estimates (Fig. S7B), suggesting that tau expression is not a major limiting factor in the spread of tau pathology.

**Figure 5.**
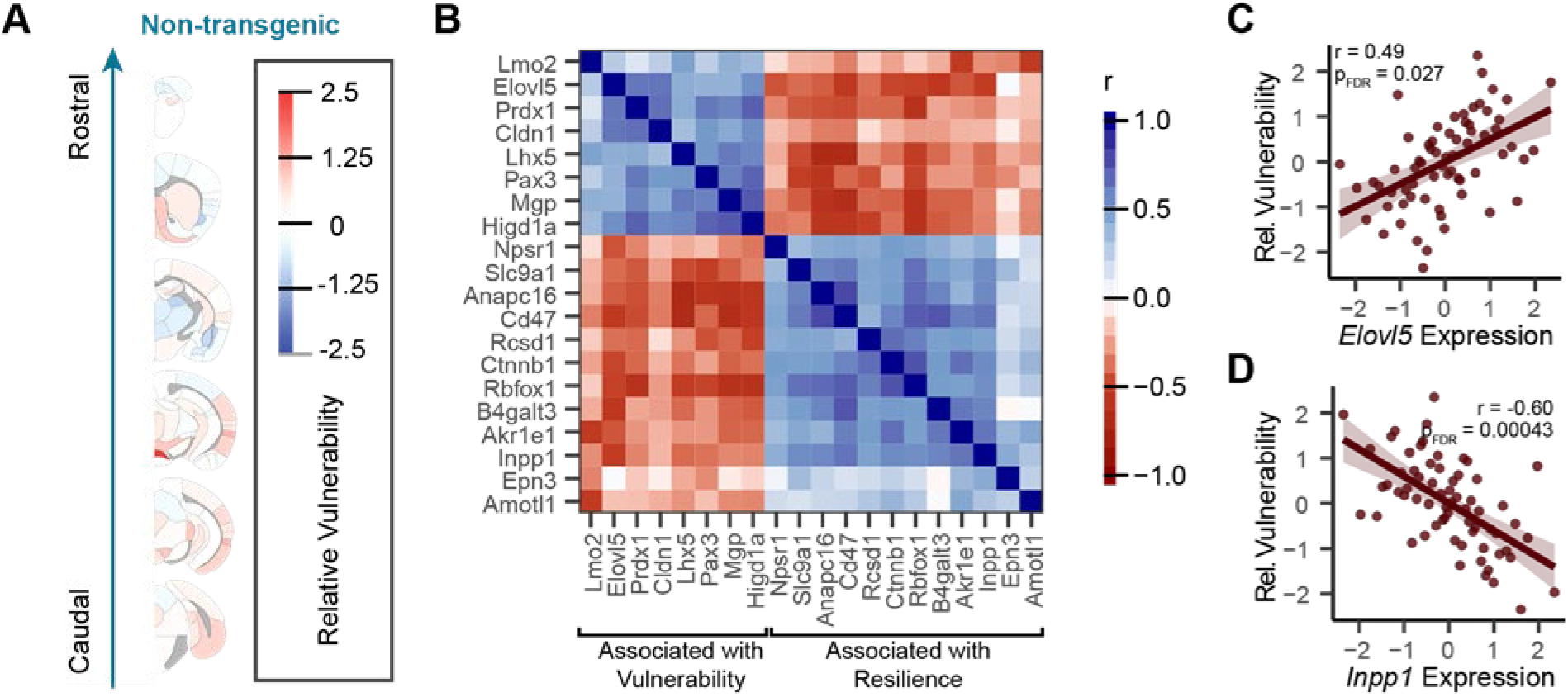
Model residuals as a predictor of regional vulnerability. **(A)** Residuals between actual tau pathology levels and pathology levels predicted by the bidirectional model w≡e averaged for each region over time and across hemisphere to give an average regional vulnerability to tau pathology. Here, those values are plotted as an anatomical heat map. (B) Spatial similarity of expression patterns between regional vulnerability-related genes (FDR corrected p<0.05 cut off for inclusion). Gene expression patterns fall into two groups of genes that are positively and negatively correlated with regional vulnerability estimates, and thereby associated with regional vulnerability and resilience, respectively. (C) Normalized relative regional vulnerability is plotted as a function of normalized *Elovol5* expression. (D) Normalized relative regional vulnerability is plotted as a function of normalized *Inppl* expression. For panels (C) and (D) the solid line represents the line of best fit and the shaded ribbons represent the 95% prediction intervals. The r and p values for the Pearson correlation between vulnerability and gene expression are noted on the plots.Plotting the gene expression versus vulnerability of individual regions (Fig. 5C, 5D, S7C) demonstrates that this association is not driven by a few regions but is spread across brain regions. In the future, gene expression patterns may be useful as predictors of regional vulnerability in computational models of pathology spread.

We sought to further investigate gene expression patterns associated with regional vulnerability by performing a genome-wide search for genes whose expression patterns measured by the ABA were spatially similar to our regional vulnerability measure. Many gene expression patterns correlated with regional vulnerability. After adjusting false discovery rate to *q* < 0.05 ^33^, we found 20 genes had expression patterns with a statistically significant spatial correlation with relative regional vulnerability to tau pathology (Fig. 5B). Twelve of these genes showed expression patterns that were negatively associated with regional vulnerability, and eight of these genes showed expression patterns that were positively associated with vulnerability.

### *In silico* seeding from alternate sites

In addition to inferring mechanisms of spread through network modeling, we can also extend the value of our validated network models by generating predictions of tau spread patterns from alternate injection sites, assuming that the rates of spread and contributions of anterograde versus retrograde spread are the same for those injection sites. For example, one of the earliest sites with tau pathology outside of the brainstem in humans is the entorhinal cortex ^34^. However, this site is difficult to inject reproducibly due to it lateral location. *In silico* modeling of tau pathology spread from this injection site shows a more lateralized spreading pattern that largely affects hippocampal and parahippocampal regions, with spread to contralateral regions occurring relatively late (Fig. 6A). We chose to also model a caudoputamen injection site (Fig. 6B) to compare tau pathology spread to previously published data modeling α-synuclein pathology spread. While tau pathology is predicted to spread to some conserved regions, including the substantia nigra and frontal cortical regions, there is also more engagement of thalamic and mesencephalic regions and less engagement of contralateral regions by tau than with α-synuclein pathology.

**Figure 6.**
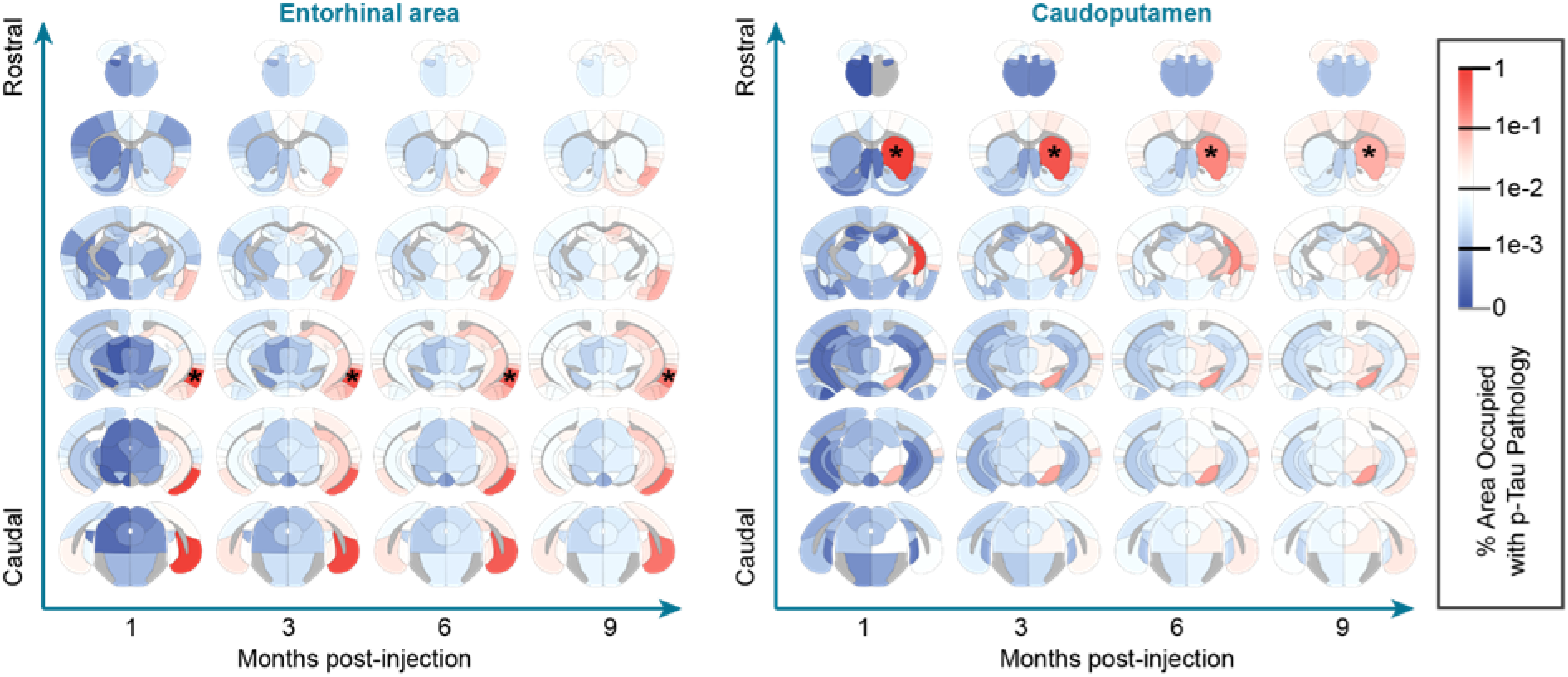
*In silico* seeding from alternate sites. Using the time diftusivity constant estimated by model fitting to empirical tau pathology spread, we estimated the distinct spreading patterns that arise after injection into alternate sites in the entorhinal area (A) and caudoputamen (B). Estimated spread is plotted as a heat map in anatomical space with blue indicating regions with minimal estimated pathology and red indicating regions with elevated pathology. * Sites of injections.

### G2019S LRRK2 mice have altered tau pathology patterns

To assess alterations in tau pathology distribution and spread related to LRRK2, we performed quantitative analysis of pathology in LRRK2^G2019S^ mice. We first established that in the absence of pathological tau injection, LRRK2^G2019S^ mice do not accumulate detectable tau pathology up to 12 months of age (Fig. S8). Following pathological tau injection, LRRK2^G2019S^ mice accumulate tau pathology in similar regions and over a similar time period as NTG mice (Fig. 7A), suggesting that overall spreading is constrained by anatomical connectivity. However, there are clear differences in the regional distribution of tau pathology in LRRK2^G2019S^ mice (Fig. 7B). There exists no overall increase or decrease in pathology in LRRK2^G2019S^ mice, suggesting that LRRK2 is not functioning to simply phosphorylate tau pathology or decrease its degradation, but likely functions to alter tau pathology spread. In some regions, like the injected iDG and highly-connected iSUM, tau pathology is almost identical in NTG and LRRK2^G2019S^ mice (Fig. 7C, 7D). In contrast, other regions, especially those which require extended periods of time to exhibit pathology show elevated pathology in LRRK2^G2019S^ mice (Fig. 7C, 7D). These results suggest that LRRK2^G2019S^ is altering some aspect of tau pathology spread.

**Figure 7.**
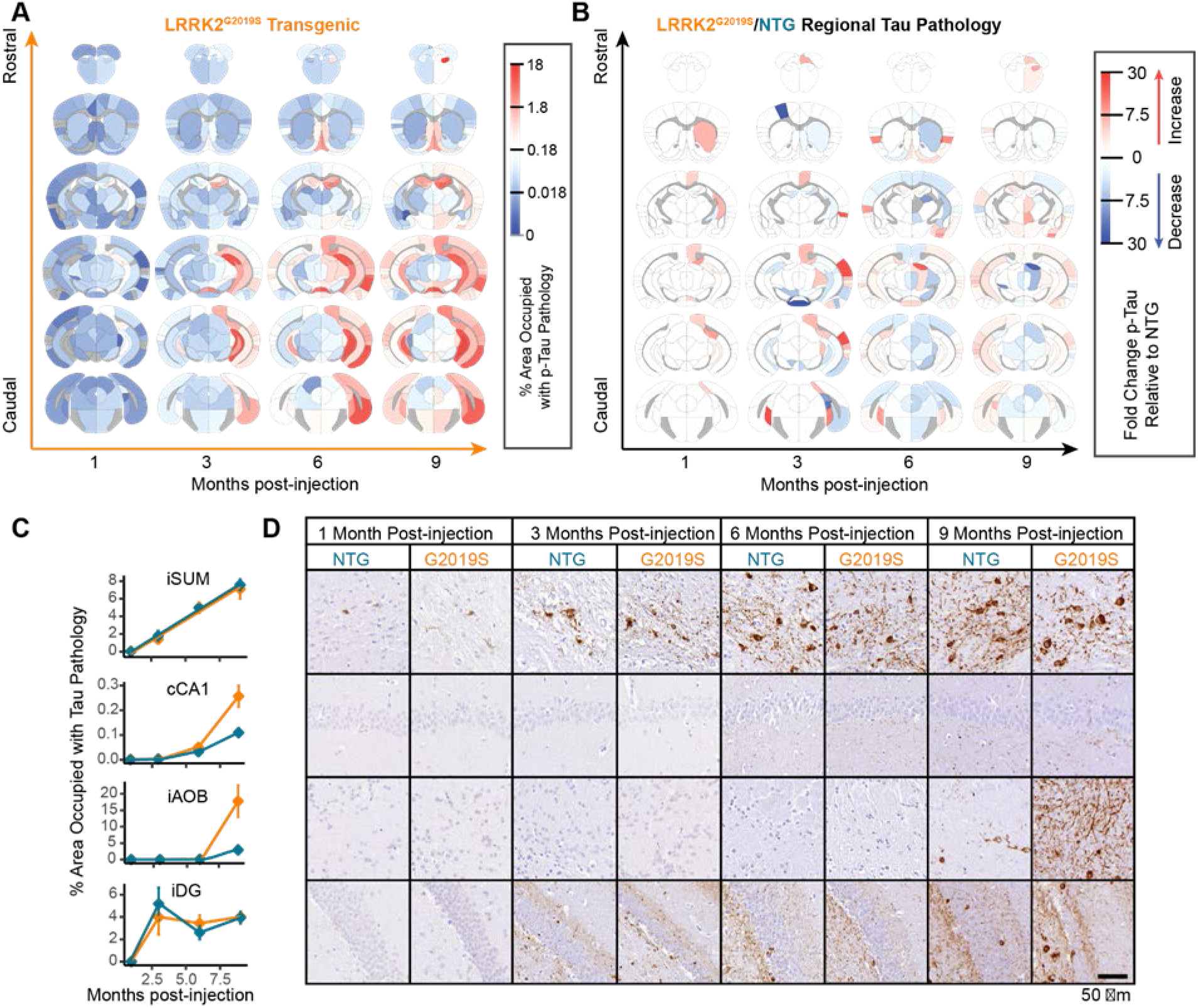
G2019S LRRK2 mice have altered tau pathology patterns. (A) Regional pathology measures for LRRK2^G2019S^ mice plotted on anatomical scaffolds as a heat map. with blue representing minimal pathology, white representing moderate pathology, and red representing substantial pathology. (B) The fold-change between NTG and LRRK2^G2O:19S^ mice is plotted on anatomical maps as a heatmap with blue representing regions with higher pathology in NTG mice, and red representing regions with higher pathology in LRRK2^G2019s^ mice. (C) The % of area occupied with tau pathology is plotted as a function of time for 4 different regions (iSUM, cCAl, iAOB, iDG) demonstrating four distinct pattern of pathology propagation in NTG and LRRK2 ^G2019s^ mice. Some regions, like the iSUM and iDG, show’ similar tau pathology progression in NTG and LRRK2^G2019S^ mice. In contrast, the cCAl and iAOB, although similar at 1 and 3 MPI show’ enhanced pathology in LRRK2^G2019S^ mice at 6 and 9 MPI. (D) Representative images of the regions plotted in panel (C) stained for p-tau and directly adjacent to the plots demonstrating the pathology pattans. Representative images for all regions can be found in the Supplementary Material. Scale bar = 50 μm.

### The G2019S LRRK2 genetic risk factor alters network dynamics of tau pathology spread

To gain a deeper understanding into what parameters of tau pathology spread may be altered, we modeled spread in LRRK2^G2019S^ mice using network diffusion. As suggested by the overall quantitative pathology pattern (Fig. 7A), tau spread can be well explained in LRRK2^G2019S^ mice by anatomical connectivity (Fig. 8A). Similar to NTG mice, tau pathology spread in LRRK2^G2019S^ mice is only moderately fit at 1 MPI, but shows improved fit at 3, 6, and 9 MPI, suggesting that anatomical connectivity is also a major factor driving tau pathology spread in LRRK2^G2019S^ mice. We next sought to explain the regional differences in pathology in LRRK2^G2019S^ mice. We first assessed the relationship of estimated regional vulnerability in NTG mice to the difference in pathology in LRRK2^G2019S^ mice (Fig. 8B). Interestingly, there is a negative correlation between NTG vulnerability and the difference in pathology at 1 and 3 MPI, suggesting that at those timepoints, regions that are resilient in NTG mice show elevated pathology in LRRK2^G2019S^ mice, whereas regions that are already vulnerable do not show a further elevation in pathology in LRRK2^G2019S^ mice. At 6 and 9 MPI, the relationship between NTG vulnerability and pathology difference is diminished.

**Figure 8.**
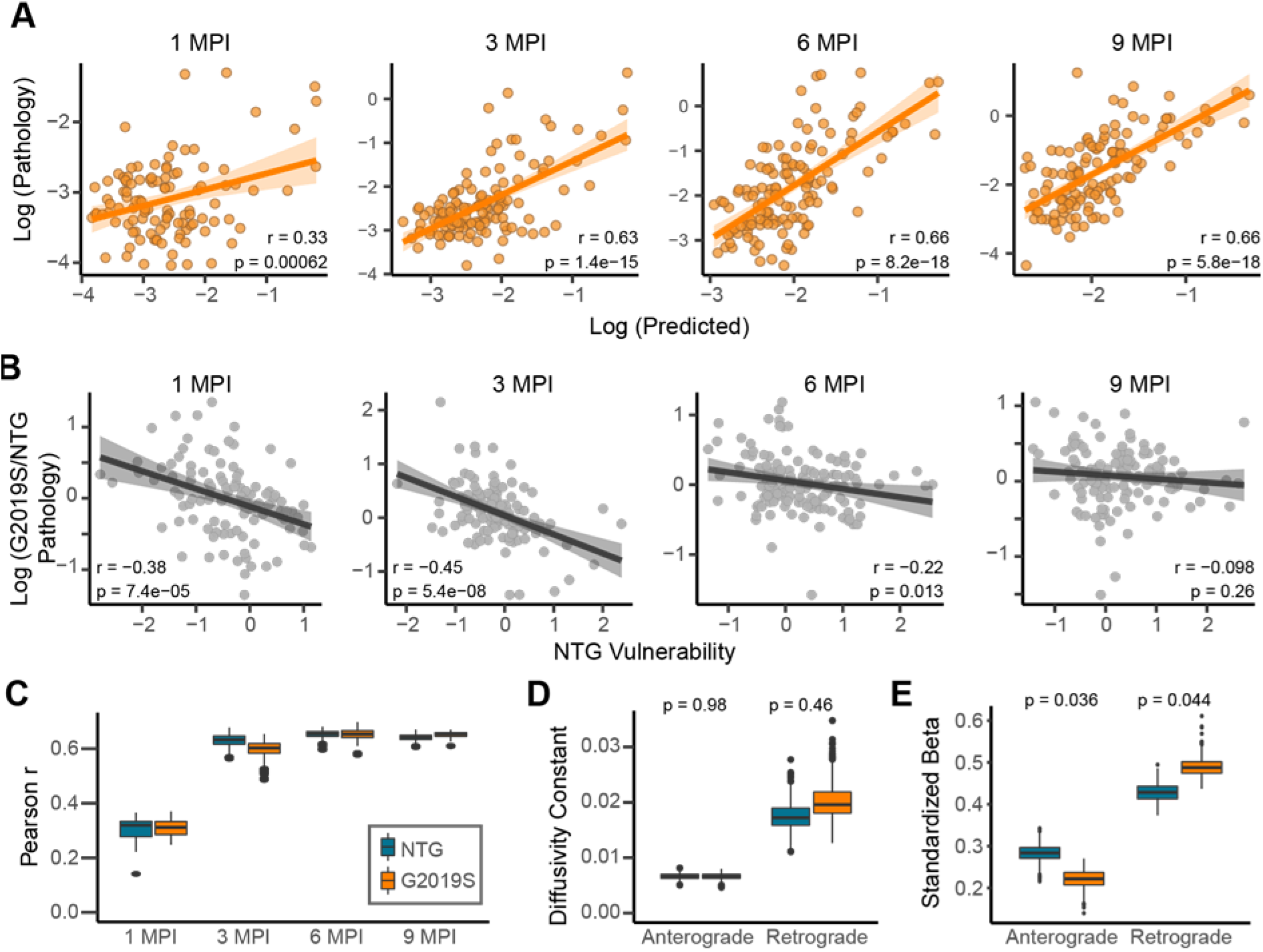
The G2019S LRRK2 genetic risk factor alters network dynamics of tau pathology spread. (A) (A) Predictions of regional log tau pathology (x-axis) in LRRK2^G2019S^ mice from spread models based on retrograde and anterograde anatomical connections, plotted against log actual regional tau pathology values (y-axis) at 1. 3,6, and 9 MPI. Solid lines represent the lines of best fit, and the shaded ribbons represent the 95% prediction intervals. The r and p values for the Pearson correlation between model fitted values and observed pathology are noted on the plots. (B) Plots of the NTG vulnerability measure versus log G2019S/NTG pathology, showing a negative correlation between the two measures especially at early time points (1 and 3 MPI) which levels off at later time point (6 and 9 MPI). Solid lines represent the lines of best fit, and the shaded ribbons represent the 95% prediction intervals. The r and p values for the Pearson correlation between G2019S/NTG pathology ratio and NTG vulnerability are noted on the plots. (C) Distributions of model fit (Pearson r) for fitting data to bootstrap samples of mice. NTG and G2O do not differ in model fit (non-parametric, two-tailed test). (D) Distributions of diffusivity constants reveals greater inter-sample variability in retrograde constants compared to anterograde. LRRK2^G2019S^ and NTG do not differ in time *constants*. (E) Anterograde and retrograde betas differ between NTG and LRRK2^G2019S^ mice, with LRRK2^G2019s^ preferentially spreading in the retrograde direction.

We next sought to determine whether the difference in regional pathology distribution in LRRK2^G2019S^ mice is related to a difference in spread along anterograde or retrograde connections. In order to infer the mechanisms of network spread affected by the LRRK2^G2019S^ mutation, we fit the bidirectional model on bootstrapped samples of NTG and LRRK2^G2019S^ mice in order to obtain distributions of model parameters; namely, the diffusivity constants and regression weights that measure the relative importance of anterograde and retrograde spread. We observed no significant difference in the overall fit of NTG and LRRK2^G2019S^ mice over time (Fig. 8C), suggesting that our network model adequately captures tau spread in LRRK2^G2019S^ mice. Additionally, the overall diffusivity constant along anterograde connections was no different than that along retrograde (Fig. 8D), suggesting that the rates of anterograde and retrograde spread are similar. Interestingly, the contribution of anterograde and retrograde connections over time did show differences (Fig. 8E), such that anterograde spread was less important and retrograde spread more important for explaining tau spread patterns in LRRK2^G2019S^ mice. These findings suggest that the LRRK2^G2019S^ mutation may lead to increased shunting of misfolded tau into retrograde pathways, which partly explains the differences in pathology patterns observed in these mice.

## DISCUSSION

Neurodegenerative diseases progress symptomatically as pathologies spread to previously unaffected regions of the brain. Identification of the neuropathological proteins aggregated in these diseases and subsequent staging studies demonstrated that symptom progression is associated with the presence of aggregated proteins in more and more regions ^34^. The stages of disease suggest that either there is a fine gradient of regional vulnerability such that regions become sequentially affected, or that pathology can spread through the brain by transcellular means. The latter hypothesis is more parsimonious, and mounting evidence has supported this hypothesis in recent years. The current study demonstrates for the first time using an interdisciplinary approach bridging quantitative pathology and network analysis that tau pathology patterns can be predicted by linear diffusion through the anatomical connectome with modulations of that spread by regional vulnerability.

Our study is congruous with previous work using computational modeling to understand the distribution of tau pathology and related regional atrophy in tauopathies. These studies assessed the utility of computational models to predict semi-quantitative regional tau pathology scores in transgenic mice ^35^, global atrophy MRI in AD and frontotemporal dementia ^36,37^, tau PET signal in AD ^38^, or general histopathological patterns of AD ^39^. While each of these studies employed different modeling parameters, each found that a model incorporating spread along anatomical connections was the most efficient and accurate predictor of tau pathology or related atrophy patterns. Our study has now extended these previous efforts in three important ways. First, the current study utilized tau seeding in a non-transgenic mouse, giving our model a precise spatiotemporal starting point, as defined by the site and time of pathogenic tau injection. Second, we utilized quantitative measures of mouse tau pathology in 194 regions of the mouse brain, providing several log-fold depth of data throughout the brain. Third, mice were sacrificed at 4 time points, following injection of pathogenic tau, providing pseudo-longitudinal data for model fitting. Our modeling of this data is consistent with previous studies, finding that tau pathology spreads in a pattern that is well-described by anatomical connectivity. Further, we were able to delineate the contribution of anterograde and retrograde connections to this spread through a rigorous model comparison approach, assess the kinetics of spread, understand regional vulnerability to pathology, compare regional vulnerability to regional gene expression, and assess the impact of a genetic risk factor on the spread of tau pathology.

This study has several limitations. One is the resolution of tau pathology data and the reliance on mesoscale connectivity measures to model tau pathology spread. Tau pathology was assessed at a mesoscale to match the mesoscale connectivity atlas generated by the Allen Institute ^31^. However, this scale obscures more granular information regarding, for example, the cortical laminar distribution of pathology. Higher-resolution connectivity maps are currently being optimized, and future studies would benefit from higher-resolution pathology maps to match the higher resolution connectivity maps. A second limitation is the use of a transgenic mouse overexpressing LRRK2. When this study began, this model was desirable because LRRK2 is expressed on the endogenous mouse promoter, ensuring that its regional expression matched that of endogenous LRRK2. However, next-generation models with a knock-in of the LRRK2 mutation are now available ^40–42^, and future work should validate the effects of LRRK2 on tau spread in these models.

Despite recent efforts, the neuropathological substrate of *LRRK2* PD has been elusive. While LRRK2 mutation carriers all have degeneration of substantia nigra neurons, many of them do not accrumulate α-synuclein Lewy bodies in the brain ^11,12^. This fact has left some question as to the neuropathological substrate of degeneration in these patients. In the search for an alternate pathology, it has now been recognized that most LRRK2 mutation carriers have tau pathology to varying degrees ^11,13,14^ Although it is not clear that tau pathology is responsible for dopaminergic neuron death in *LRRK2* mutation carriers, it does raise the possibility that mutations in *LRRK2* can predispose patients to developing tau pathology.

Interestingly, hyperphosphorylated tau is a well-cataloged feature present in LRRK2 mutant mice ^43,44^, flies ^45^, and iPSC-derived neurons ^46^. It has also been demonstrated that LRRK2 mutations can lead to phosphorylation of tau ^47–50^. Although it is not clear that this hyperphosphorylated tau is pathological, it is possible that it represents a pool of tau that is more rapidly recruited upon the introduction of a pathogenic tau seed. The current study was conducted in mice up to 12 months of age, and the LRRK2^G2019S^ mice in this study showed no evidence of phosphorylated tau accumulation without pathogenic tau injection. This observation suggests that the changes in tau pathology spread are likely due not to phosphorylation, but rather to alterations in cellular mechanisms such as protein release, internalization, transport, or degradation. This hypothesis is supported by another recent study showing enhanced neuronal transmission of virally-expressed tau ^51^. Other studies found that overall tau pathology was not affected in transgenic mice which exhibit broad tau pathology controlled by the transgene promoter ^51,52^.

One possible cellular mechanism regulating the change in tau pathology spread in LRRK2^G2019S^ mice is elevated presynaptic vesicle release, a phenomenon which has been observed in LRRK2^G2019S^ mice ^42^. This enhanced release could provide an increased opportunity for extracellular tau to be internalized by recipient neurons and thereby enhance retrograde transmission of pathology. Previous research has demonstrated that tau is readily released by neurons ^22–24^ in proportion to neuronal activity ^25,26^. Future research incorporating functional connectivity strength and tau receptor distribution as regulators of tau spread may help clarify how tau pathology spread is regulated.

In conclusion, the current study demonstrates that tau pathology spreads from an initial injection site through the brain via neuroanatomical connectivity. This spread can be modulated by a genetic risk factor for PD. Future work should substantiate that this alteration is related to LRRK2 kinase activity and explore whether LRRK2 inhibitors would be a viable therapeutic treatment for tauopathies.

## ACKNOWLEDGEMENTS

We would like to thank members of the laboratory for their feedback in developing this manuscript. This study was supported by NIH grants: T32-AG000255 (V.M.Y.L), P30-AG10124 (J.Q.T.) U19-AG062418 (J.Q.T.), P50-NS053488 (J.Q.T.), R01-NS099348 (D.S.B.), F30 MH118871-01 (E.J.C); NSF grants PHY-1554488 (D.S.B) and BCS-1631550 (D.S.B); and a Michael J. Fox Foundation grant 16879 (M.X.H.). D.S.B. also acknowledges support from the John D. and Catherine T. MacArthur Foundation, the ISI Foundation, the Alfred P. Sloan Foundation, and the Paul G. Allen Foundation.

## AUTHOR CONTRIBUTIONS

Conceptualization, M.X.H., V.M.Y.L.; Methodology, M.X.H., E.J.C., V.M.Y.L; Software, E.J.C., D.S.B.; Formal Analysis, M.X.H., E.J.C.; Investigation, M.X.H., E.J.C., H.L.L., L.C., B.Z., H.J.B., R.J.G., M.F.O.; Resources, L.C.; Writing-Original Draft, M.X.H., E.J.C.; Writing-Review and Editing, All; Visualization, M.X.H., E.J.C.; Supervision, M.X.H., D.S.B., J.Q.T., V.M.Y.L.; Funding Acquisition, M.X.H., D.S.B., J.Q.T., V.M.Y.L.

## MATERIALS AND METHODS

### Mice

All housing, breeding, and procedures were performed according to the NIH Guide for the Care and Use of Experimental Animals and approved by the University of Pennsylvania Institutional Animal Care and Use Committee. C57BL/6J (NTG, JAX 000664, RRID: IMSR_JAX:000664) and B6.Cg-Tg(Lrrk2*G2019S)2Yue/J (G2019S, JAX 012467, RRID: IMSR_JAX:012467) mice have been previously described ^16^. The current G2019S BAC line was backcrossed to C57BL/6J mice for >10 generations and bred to homozygosity at loci as determined by quantitative PCR and outbreeding. The expression level of G2019S LRRK2 was thereby stabilized in this line of mice. All experiments shown use homozygous G2019S mice. Both male and female mice were used for this study. For *in vivo* experiments, both male (n=24) and female (n=19) mice were used and were 3-4 months old at the time of injection. No influence of sex was identified in the measures reported in this study.

### Primary Hippocampal or Neuron Cultures

Primary hippocampal neuron cultures were prepared as previously described ^53^ from postnatal day (P) 1 non-transgenic or LRRK2^G2019S^ transgenic mice. Dissociated hippocampal or cortical neurons were plated at 17,000 cells/well (96-well plate) or 1,000,000 cells/well (6-well plate) in neuron media (Neurobasal medium (ThermoFisher 21103049) supplemented with B27 (ThermoFisher 17504044), 2 mM GlutaMax (ThermoFisher 35050061), and 100 U/mL penicillin/ streptomycin (ThermoFisher 15140122).

### Recombinant Tau PFFs

Purification of recombinant human tau and generation of tau X-T40 PFFs was conducted as described elsewhere 15. The plasmid containing the gene of interest was transformed into BL21 (DE3) RIL-competent *E. coli* (Agilent Technologies Cat#230245). A single colony from this transformation was expanded and tau protein was purified by cationic exchange using fast protein liquid chromatography as previously described ^54^.

Tau fibrillization was induced in the absence of co-factors by incubation of 40 μM recombinant T40 (4R2N tau) monomer with 2 mM DTT in Dulbecco’s phosphate-buffered saline (DPBS, Corning Cat#21-031-CV) and incubated shaking at 1,000 rpm and at 37°C for 7 days to create passage 1 of fibrils. A second passage was set up by incubating 10% of the first passage with 90% fresh tau monomer at a final concentration of 40 μM tau. This passaging was repeated until tau was recovered in the pellet fraction in a stochastic subset of the reactions. Conversion to PFFs was validated by sedimentation at 100,000 x *g* for 30 minutes at 22°C. Equal volumes of supernatant and pellet fraction were loaded and analyzed by Coomassie blue staining of SDS-PAGE gels.

### Human Tissue

All procedures were done in accordance with local institutional review board guidelines of the University of Pennsylvania. Written informed consent for autopsy and analysis of tissue sample data was obtained either from patients themselves or their next of kin. Cases used for extraction (Fig. S1) of PHF tau were selected based upon a high burden of tau pathology by immunohistochemical staining. Cases used for extraction were balanced by sex (female = 2; male = 3) and were frozen an average of 6 hours post-mortem. Differences in sex were not assessed because these cases were only utilized for protein extraction.

### LRRK2 Inhibitor Treatments

LRRK2 inhibitors PF-475 and PF-360 were synthesized at Pfizer, Inc. MLi-2 was obtained from Tocris Bioscience (Cat#5756). All LRRK2 inhibitors were reconstituted at 10 mM in DMSO and stored at −20°C. They were further diluted to the final concentration indicated in neuron media with DMSO as a vehicle control.

### Tau XT-40 PFF/AD PHF Treatments

#### Primary Neurons

For treatment of neurons, X-T40 PFF tau and AD PHF tau were vortexed and diluted with Dulbecco’s phosphate-buffered saline (DPBS, Corning Cat#21-031-CV). They were then sonicated on high for 10 cycles of 30 seconds on, followed by 30 seconds off (Diagenode Biorupter UCD-300 bath sonicator). Tau was then diluted in neuron media to the noted concentrations and added to neuron cultures at 7 DIV. Neuron cultures were harvested 14-21 days post-treatment (DPT), as noted.

#### Mice

All surgery experiments were performed in accordance with protocols approved by the Institutional Animal Care and Use Committee (IACUC) of the University of Pennsylvania. AD PHF tau was vortexed and diluted with DPBS to 0.4 mg/mL. Tau was sonicated (QSonica Microson™ XL-2000; 60 pulses; setting 1.5; 1 sec/pulse). Mice were injected when 3-4 months old. Mice were deeply anaesthetized with ketamine/xylazine/acepromazine and injected unilaterally by insertion of a single needle into the right forebrain (coordinates: −2.5 mm relative to Bregma, +2.0 mm from midline) targeting the hippocampus (2.4 mm beneath the skull) with 1 μg tau (2.5 μL). Needle was then retracted to 1.4 mm beneath the skull, targeting the overlaying cortex and another 1 μg tau (2.5 μL) was injected. Injections were performed using a 25 μL syringe (Hamilton, NV) at a rate of 0.4 μL/minute. After 1 to 9 months, mice were perfused transcardially with PBS, brains were removed and underwent overnight fixation in 70% ethanol in 150 mM NaCl, pH 7.4.

### Human Brain Sequential Detergent Fractionation

Frozen postmortem human frontal or temporal cortex brain tissue containing abundant tau-positive inclusions was selected for sequential extraction based on IHC examination of these samples as described ^55^ using previously established methods. These brains were sequentially extracted with increasing detergent strength as previously described 15. After thawing, meninges were removed and gray matter was carefully separated from white matter. Gray matter was weighed and suspended in nine volumes (w/v) high salt (HS) buffer (10 mM Tris-HCL (pH 7.4), 800 mM NaCl, 1 mM EDTA, 2 mM dithiothreitol [DTT], protease and phosphatase inhibitors and PMSF) with 0.1% sarkosyl and 10% sucrose, followed by homogenization with a dounce homogenizer and centrifugation at 10,000 x *g* for 10 minutes at 4°C. The resulting pellet was re-extracted with the same buffer conditions and the supernatants from all extractions were filtered and pooled.

Additional sarkosyl was added to the pooled supernatant to reach a final concentration of 1% and the supernatant was nutated for 1 hour at room temperature. The samples were then centrifuged at 300,000 x *g* for 60 minutes at 4°C. The pellet, which contains pathological tau, was washed once with PBS and resuspended in 100 μL of PBS per gram of gray matter by passing through a 27G/0.5 in. needle. The pellets were further suspended by brief sonication (QSonica Microson™ XL-2000; 20 pulses; setting 2; 0.5 sec/pulse). The suspension was centrifuged at 100,000 x *g* for 30 minutes at 4°C. The pellet was suspended in one-fifth to one-half the pre-centrifugation volume, sonicated briefly (60-120 pulses; setting 2; 0.5 sec/pulse) and centrifuged at 10,000 x *g* for 30 minutes at 4°C. The final supernatant was utilized for all studies and is referred to as AD PHF tau. All extractions were characterized by Western blotting, sandwich ELISA for tau, α-synuclein and Aβ 1–40, Aβ 1–42, and validated by immunocytochemistry in primary neurons from non-transgenic mice. For the extractions used in this study, tau constituted 16.1-35.7% of the total protein, while α-synuclein and Aβ constituted 0.011% or less of total protein.

### Immunoblotting

Total protein concentration in each sample was determined by a bicinchoninic acid colorimetric assay (Fisher Cat#23223 and 23224), using bovine serum albumin as a standard (Thermo Fisher Cat#23210). Protein was resolved on 5-20% gradient polyacrylamide gels using equal protein loading. Proteins were transferred to 0.2 um nitrocellulose or PVDF membranes and detected with primary antibodies targeting LRRK2 (ab133474, Abcam, RRID: AB_2713963, 1:500), pS935 LRRK2 (ab133450, Abcam, RRID:AB_2732035, 1:400) or GAPDH (2-RGM2, Advanced Immunological, RRID:AB_2721282, 1:5000). Primary antibodies were detected using IRDye 800 (Li-cor 925-32210) or IRDye 680 (Li-cor 925-68071) secondary antibodies, scanned on Li-cor Odyssey Imaging System and analyzed using Image Studio software. LRRK2 and pS935 LRRK2 values were normalized to GAPDH as an internal loading control, then further normalized to the mean of all control samples.

### Immunocytochemistry

Primary neurons treated with X-T40 tau PFFs were fixed with 4% paraformaldehyde, 4% sucrose in phosphate-buffered saline and washed five times in PBS. Cells were permeabilized in 3% BSA + 0.3% TX-100 in PBS for 15 minutes at room temperature. After a PBS wash, cells were blocked for 50 minutes with 3% BSA in PBS prior to incubation with primary antibodies for 2 hours at room temperature. Primary antibodies used were targeting pS202/T205 tau (AT8, ThermoFisher Cat#MN1020, 1:1000), MAP2 (17028, CNDR, 1:2000). Cells were washed 5x with PBS and incubated with secondary antibodies for 1 hour at room temperature. After 5x wash with PBS, cells were incubated in DAPI (ThermoFisher Cat#D21490, 1:10,000) in PBS. Primary neurons treated with AD PHF tau were stained differently due to the possible presence of p-tau signal in the human-derived material added to cultures. Soluble protein was extract with 2% HDTA for 10 minutes at room temperature. Neurons were then fixed and stained as above, except that a primary antibody that selectively binds mouse tau (T49, CNDR, 1:2500) was utilized to detect neuronal tau inclusions.

96-well plates were imaged on an In Cell Analyzer 2200 (GE Healthcare) and analyzed in the accompanying software. A standard intensity-based threshold was applied to MAP2 and p-tau or mouse tau channels equally across plates and the positive area was quantified. All quantification was optimized and applied equally across all conditions.

### Immunohistochemistry

After perfusion and fixation, brains were embedded in paraffin blocks, cut into 6 μm sections and mounted on glass slides. Slides were then stained using standard immunohistochemistry as described below. Slides were de-paraffinized with 2 sequential 5-minute washes in xylenes, followed by 1-minute washes in a descending series of ethanols: 100%, 100%, 95%, 80%, 70%. Slides were then incubated in deionized water for one minute prior to antigen retrieval as noted. After antigen retrieval, slides were incubated in 7.5% hydrogen peroxide in water to quench endogenous peroxidase activity. Slides were washed for 10 minutes in running tap water, 5 minutes in 0.1 M tris, then blocked in 0.1 M tris/2% fetal bovine serum (FBS). Slides were incubated in primary antibody overnight. For pathologically-phosphorylated tau, pS202/T205 tau (AT8, ThermoFisher Cat#MN1020) was used at 1:10,000 with microwave antigen retrieval.

Primary antibody was rinsed off with 0.1 M tris for 5 minutes, then incubated with goat anti-rabbit (Vector BA1000) or horse anti-mouse (Vector BA2000) biotinylated IgG in 0.1 M tris/2% FBS 1:1000 for 1 hour. Biotinylated antibody was rinsed off with 0.1 M tris for 5 minutes, then incubated with avidin-biotin solution (Vector PK-6100) for 1 hour. Slides were then rinsed for 5 minutes with 0.1 M tris, then developed with ImmPACT DAB peroxidase substrate (Vector SK-4105) and counterstained briefly with hematoxylin. Slides were washed in running tap water for 5 minutes, dehydrated in ascending ethanol for 1 minute each (70%, 80%, 95%, 100%, 100%), then washed twice in xylenes for 5 minutes and coverslipped in Cytoseal Mounting Media (Fisher 23-244-256).

Slides were scanned into digital format on a Lamina scanner (Perkin Elmer) at 20x magnification. Digitized slides were then used for quantitative pathology.

### Quantitative pathology

All section selection, annotation, and quantification was done blinded to treatment. For quantification of tau pathology, coronal sections were selected to closely match the following coordinates, relative to Bregma: 3.20 mm, 0.98 mm, −1.22 mm, −2.92 mm, −3.52, and −4.48 mm. The digitized images were imported into HALO software to allow annotation and quantification of the percentage area occupied by tau pathology. Standardized annotations were drawn to allow independent quantification of 194 gray matter regions throughout the brain. Each set of annotations was imported onto the desired section and modified by hand to match the designated brain regions. After annotation, the analysis scripts were applied to the brain to make sure that no non-pathology signal was detected. After annotation of all brains, analysis algorithms were applied to all stained sections, and data analysis measures for each region were recorded.

The total pathology analysis detects total signal above a minimum threshold. Specifically, the analysis included all DAB signal that was above a 0.099 optical density threshold, which was empirically determined to not include any background signal. This signal was then normalized to the total tissue area. A minimal tissue optical density of 0.02 was used to exclude any areas where tissue was split, and a tissue edge thickness of 25.2 μm was applied to exclude any edge effect staining.

### Computational models of pathological protein spread

Models of linear diffusion along white matter fibers have been used to predict the spread of misfolded α-synuclein in mice ^32^, as well as patterns of atrophy observed in various neurodegenerative diseases ^36,56^. In the present work, we extended these models to the spread of tau between anatomically connected brain regions from an injection site in the right hippocampus.

We model pathological spread of tau as a diffusion process on a directed structural brain network *G* = {*V, E*} whose nodes *V* are *N_a_* = 426 cortical and subcortical grey matter regions, and whose edges *e_ij_* ∈ *E* represent an axonal projection initiating in *V_i_* and terminating in *V_j_*, where *e_ij_* ≥ 0 for all *E*. Edge strength was quantified by the Allen Brain Institute using measures of fluorescence intensity from retrograde viral tract tracing ^31^. We define the weighted adjacency matrix of *G* as ***A*** = [***A**_ij_*], such that row *i* contains the strengths of axonal projections from region *í* to all other *N_a_* − 1 regions. We define the current levels of simulated tau pathology of all *N_a_* nodes at time *t* as the vector ***x***(*t*). We make empirical measurements of tau pathology at *t* = 1, 3, 6, and 9 months post injection (MPI) in an *N_c_* = 134 region vector ***y***(*t*), which is a spatially coarse grained version of ***x***(*t*). Note that ***y***(*t*) = *f*(**x**(*t*)), where *f* is a linear transformation that sets each element (region) of ***y***(*t*) equal to the arithmetic mean of the elements (regions) of ***x***(*t*) that lie within the regions of ***y***(*t*). This transformation is needed to avoid quantifying pathology in many of the smaller regions in the *N_a_*-dimensional space used by Allen Brain Institute ^31^, which are difficult to identify reliably across mice in practice.

In the simplest case of our models of pathological network spread, we simulated the spread of tau pathology throughout the *N_a_* anatomically connected brain regions in ***A*** to compute the predicted pathology 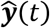 in *N_c_* empirically assessed brain regions as a function of a set of seed regions *s* ∈ *V* using the form

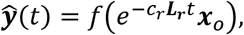

where the retrograde, out-degree graph Laplacian

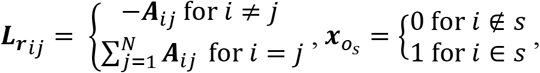

*c_r_* is a time constant representing the global speed of retrograde spread, *f* is the linear transformation described above that converts the *N_a_* region space to the *N_c_* region space, and *t* is in units of months. In this manuscript, s contains the Allen Brain Atlas regions DG, CA1, CA3, VISam, and RSPagl, in order to account for experimental variability in targeting a hippocampal injection site. Note that *c_r_* tunes the time scale of the system, which is necessary due to the fact that the units of connection strength are arbitrary relative to the units of pathology. To fit this model, we swept through values of *c_r_* from 10^-5^ to 0.2 and chose the value of *c_r_* that maximized the average Pearson correlation coefficient between log_10_***y***(*t*) and 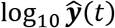 over *t* = [1 3 6 9]. Empirically, the value of this correlation plateaued for values of *c_r_* larger than 0.2, justifying this upper bound on *c_r_*. Note that ***L*** is the out-degree Laplacian, a version of the well-characterized graph Laplacian designed for directed graphs ^57^. Intuitively, this model posits that pathology spreads retrogradely from region *i* to other regions at a rate proportional to the number of synapses projecting onto *i* from those regions, while pathology at region *i* decays as a function of the sum of the strength of projections into region *i*.

However, recent studies by this group ^32^ and others ^58^ has suggested that both anterograde and retrograde spread of pathology contribute to neurodegenerative disease progression. Thus, we expanded the retrograde model described above to a bidirectional model including anterograde spread, using the form

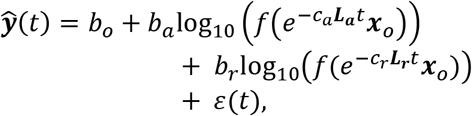

where the anterograde, out-degree graph Laplacian 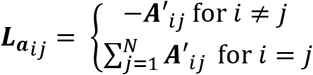, *c_a_* is a time constant representing the global speed of anterograde spread, *b_o_* is an intercept, *b_a_* is a weight for the importance of anterograde spread, *b_r_* is a weight for the importance of retrograde spread, *t* is time, and *ε* is an error term. We used the *optim* function in R to solve for the combination of *c_r_* and *c_a_* that maximizes the average Pearson correlation coefficient between log_10_***y***(*t*) and 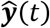 over *t* = [1 3 6 9], with a linear regression inside of the objective function to solve for *b_o_, b_a_*, and *b_r_* aggregating across all time points for each *c_r_* and *c_a_*. Importantly, the linear regression coefficients *b_a_*, and *b_r_*, when standardized, capture the relative importance of anterograde and retrograde spread at time *t*, respectively, while controlling for the potentially ambiguous overlapping contributions of the two modes of transmission.

We also defined intrinsic regional vulnerability based on *ε*(*t*), the error term in the model above. Intuitively, if *ε_i_*(*t*) is large, then this model underpredicted pathology at region *i* such that region *i* is more vulnerable to pathology than expected based upon bidirectional linear diffusion, and *vice versa* for regions with small values of *ε*(*t*). We hypothesized that both static, intrinsic regional vulnerability as well as possible time-dependent vulnerability is captured by *ε*(*t*). Therefore, we averaged *ε*(*t*) over hemispheres and over *t* = [3 6 9] in order to capture static, intrinsic regional vulnerability as an 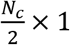 vector ***v_s_*** and average out the effects of both time-dependent vulnerability and unexplainable measurement error. We excluded 1 MPI because pathology at this time point was poorly captured by the spread model, and *ε*(1) exhibited a distinct spatial pattern of vulnerability from the patterns observed in *ε*(*t*) at *t* = [3 6 9] (Fig. S7).

#### Quantification of model specificity

In a previous study 32, we found that the substantial variance explained in misfolded α-synuclein spread by connectome-based linear diffusion models was specific to the use of iCP as the seed site *s*, which defines the vector ***x_o_***, over nearly every other region in the connectome. Here, we sought to replicate this finding in the study of tau spread. However, due to the use of multiple seed sites in the present study, additional considerations applied in ensuring a rigorous test of the model’s specificity to the experimentally motivated group of seed sites. The 5 chosen seed sites in s (DG, CA1, CA3, VISam, and RSPagl) were relatively close together (average distance of 1.96mm), so we wanted to rule out the possibility that 1) choosing multiple random sites would trivially improve model performance, and 2) selecting multiple spatially clustered random seed sites would trivially improve model performance. Thus, we choose 500 random sets of 5 seed sites *s_null_* ∈ *V*, all of which had an average distance from one another within 1.96±0.196 mm, and fit the bidirectional model detailed above using each set of distance-constrained random points *s_null_* to define the initial vector ***x_o_***. We computed a one-tailed, non-parametric *p*-value for the specificity of s by computing the percentage of times *s_null_* yielded a better fit to the observed pathology data than *s*.

#### Model comparison

The data presented in the body of the paper utilize values for the time constants, *c_r_* and *c_a_*, and time-dependent weights on retrograde and anterograde spread, *b_o_*, *b_a_*, and *b_r_*, obtained using data from all mice at every time point (***y**_full_*(*t*)). To ensure that this approach did not result in overfitting, and to rigorously test whether including both anterograde and retrograde connections improved model performance, we randomly sampled without replacement the available mice at each time point to generate ***y**_train_* (*t*) and ***y**_test_*(*t*) for each time point. These parameters were determined by applying the model fitting procedure described above on ***y**_train_*(*t*), and the model was evaluated based on its fit with ***y**_test_*(*t*) This process was repeated 500 times for models based on Euclidean distance, anterograde connections alone, retrograde connections alone, or both anterograde and retrograde connections, generating a distribution of out-of-sample fits for each time point for each model, allowing us to statistically compare the out-of-sample performance between each of the candidate models. Using a similar approach, we generated distributions of model parameters for the bidirectional model using bootstrapped samples of NTG and LRRK2 G2019S mice.

#### Network null models

To ensure that our results were specific to the retrograde spread of misfolded synuclein along neuronal processes, we repeated our analyses using several network null models. To demonstrate a general specificity of the model for the topology of the synaptic connectome represented by ***A***, we carried out a procedure that rewires the edges of *G* while exactly preserving either the out-degree or the in-degree sequence, i.e. 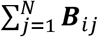 and 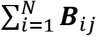, where 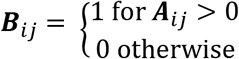. This rewiring approach tests to otherwise whether the model fit is due to a relatively basic structural property of the graph, i.e. degree, as opposed to unique, higher order topological features of the synaptic connectome. Finally, we tested the null model that spread of misfolded protein occurs simply due to diffusion through tissue based on closeness in Euclidean space. To test this model, we reconstructed ***A*** = [***A***_*ij*_] such that the edges represented the inverse Euclidean distance between region *i* and region *j*. We used the initial procedure for retrograde model fitting to test this model, because Euclidean distance is symmetric and thus the bidirectional model cannot be applied.

### Assessment of Regional Gene Expression Patterns

In order to assess the cellular and molecular characteristics of intrinsically vulnerable or resilient regions, we compared the spatial alignment between our 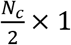 vector of regional vulnerability measurements, ***v_s_***, with microarray gene expression levels obtained from the Allen Mouse Brain Atlas (brain-map.org). After applying a previously validated quality control approach to hone in on genes with the most reliable expression measurements ^59^, we obtained ***G***, an 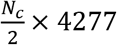 matrix of gene expression values across each brain region. Due to the non-normality of both ***v_s_*** and gene expression in ***G***, we applied a rank inverse normalization transform60 to each column of ***G*** in order to control type I error and maximize statistical power. Next, we computed 4277 spatial Pearson correlations between ***v_s_*** and each column of ***G***. We performed multiple comparisons correction by controlling the false discovery rate at *q* < 0.05.

## QUANTIFICATION AND STATISTICAL ANALYSIS

The number of samples or animals analyzed in each experiment, the statistical analysis performed, as well as the p-values for all results <0.05 are reported in the figure legends. For all *in vivo* experiments, the reported “n” represents the number of animals. For all cell culture experiments, “n” represents the number of separate cultures (e.g. one scraped or imaged well is reported as one “n”).

All cell culture data and *in vivo* non-pathological measures were analyzed in GraphPad Prism 7 using the noted statistical tests. *In vivo* pathological spread data was analyzed and all computations were performed in R (https://www.R-project.org/) ^61^ as described.

## DATA AND CODE AVAILABILITY

Primary data and code used to generate the spread modeling is available on GitHub (https://github.com/ejcorn/tau-spread).

## Supplementary Information

**Figure S1.**
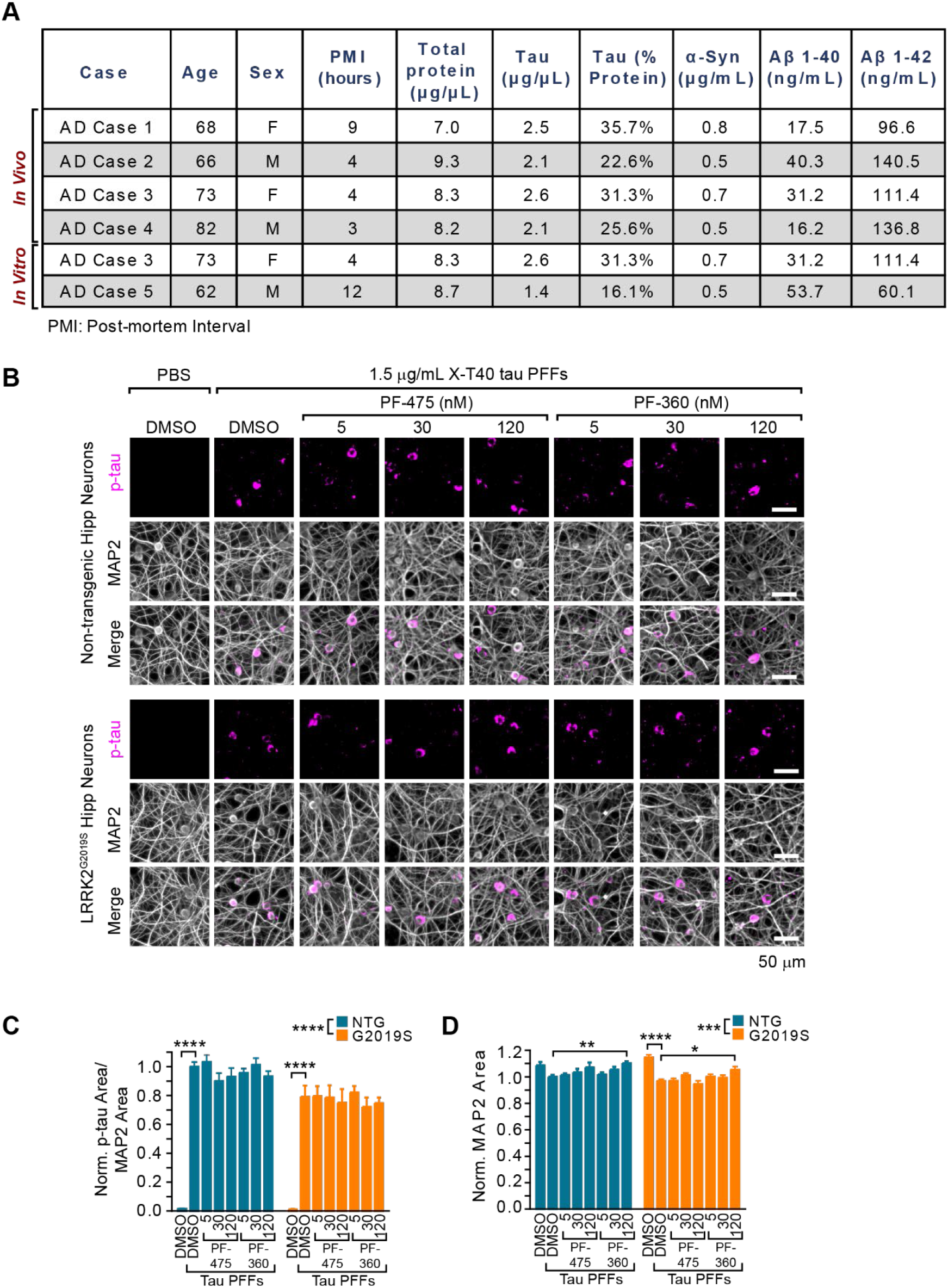
Characterization of AD PHF preparations and X-T40 primary neuron assays, Related to Figure 1. (A) Descriptive information related to the cases of AD PHFs tau prepared for these studies. Note that 5 cases were extracted. AD Case 3 was utilized for both *in vitro* and *in vivo* experiments. (B) Primary hippocampal neurons from NTG or LRRK2*G2019S mice were treated with vehicle or LRRK2 inhibitors at the noted concentrations at 5 DIV. They were further treated with X-T40 tau PFFs at 1.5 μg/mL at 7 DIV and fixed and stained for pS202/T205 tau (AT8, magenta) and MAP2 (gray) at 21 DIV. Scale bar = 50 μm. (C) Quantification of the pS202/T205 tau area normalized to MAP2 area and further normalized to NTG-DMSO-PFF. LRRK2*G2019S neurons showed a small, genotype-level significant reduction in tau pathology, and each genotype showed no tau pathology without addition of X-T40 tau PFFs (Two-way ANOVA; genotype effect ****p<0.0001, Dunnett’s multiple comparison test within genotype: NTG: DMSO-PFF vs. DMSO****p<0.0001; G2019S: DMSO-PFF vs. DMSO ****p<0.0001; All other values were not statistically significant). (D) Quantification of the MAP2 also showed a small genotype-level significant change as well as a reduction related to PFF treatment. Interestingly, the highest dose of PF-360 also elevated MAP2 area slightly (Two-way ANOVA; genotype effect ***p=0.001, Dunnett’s multiple comparison test within genotype: NTG: DMSO-PFF vs. 120 nM PF-360-PFF **p=0.0027; G2019S: DMSO-PFF vs. DMSO ****p<0.0001; DMSO-PFF vs. 120 nM PF-360 *p= 0.0216; All other values were not statistically significant). Data are represented as mean ± SEM.

**Figure S2.**
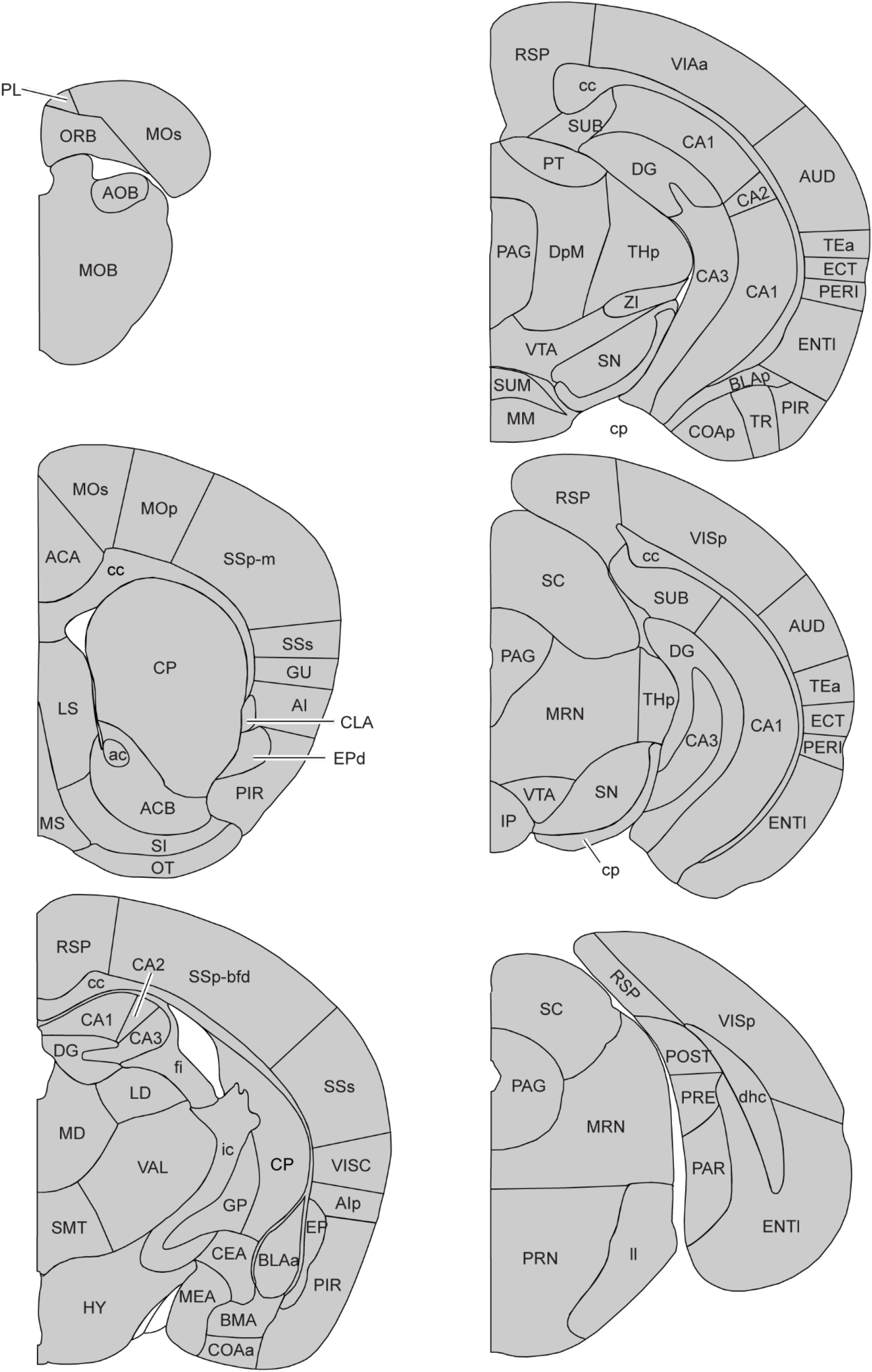
Brain region annotation key, Related to Figure 2. Each of 194 gray matter brain regions used in analysis of pathological burden are labelled here.

**Figure S3.**
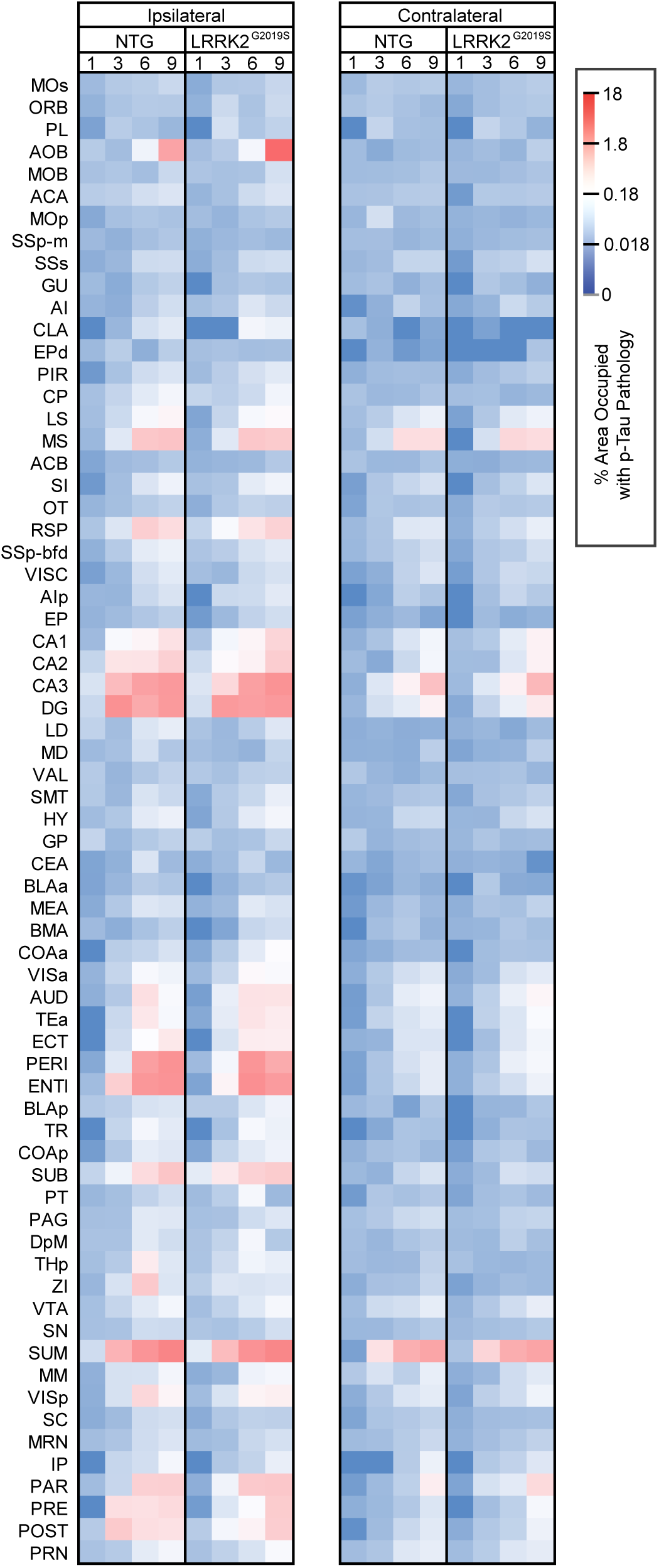
Heatmap of tau pathology in NTG and LRRK2^G2019S^ mice, Related to Figure 3. Percentage area occupied with tau pathology was measured across 194 annotations. Regions that were represented by more than one annotation in multiples sections were averaged to give a combined 134 regions. The average tau pathology in each of those regions at 1, 3, 6, and 9 MPI are displayed here as a heat map.

**Figure S4.**
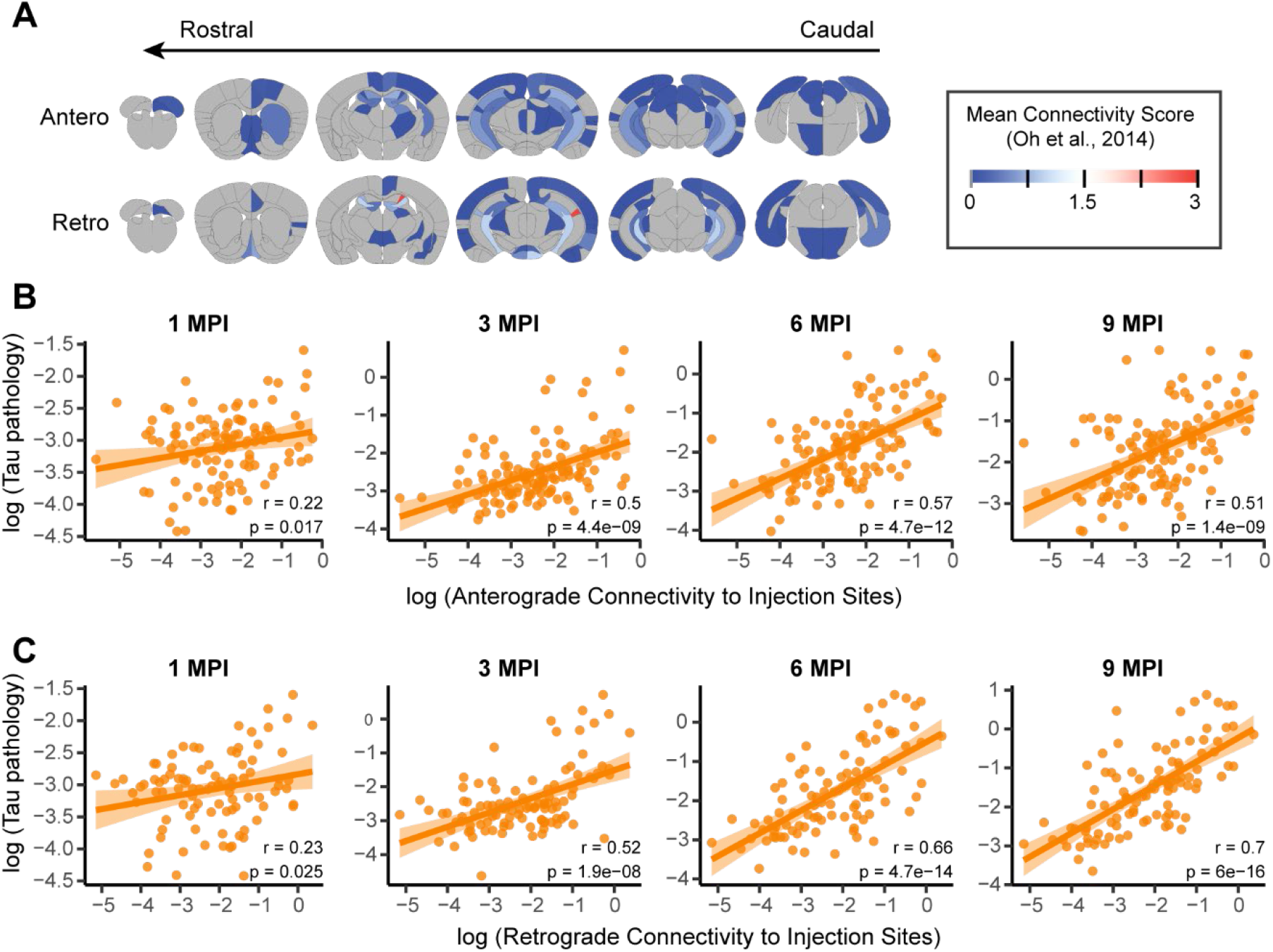
anatomical connectivity can explain spread of tau pathology, Related to Figure 4. (A) Direct anatomical connectivity was estimated for the 5 regions at the sites of injection based. There is both anterograde and retrograde connectivity largely between hippocampal and septal regions, with additional connectivity between cortical regions. (B, C) The ability of diffusion along direct connections to the injection site to predict tau pathology is assessed for both antergrade (B) and retrograde (C) connections. Note that regions with zero connectivity or zero pathology are not considered in this assessment. The solid lines represent the lines of best fit, and the shaded ribbons represent the 95% prediction intervals. The r and p values for each linear regression are noted on the plots.

**Figure S5.**
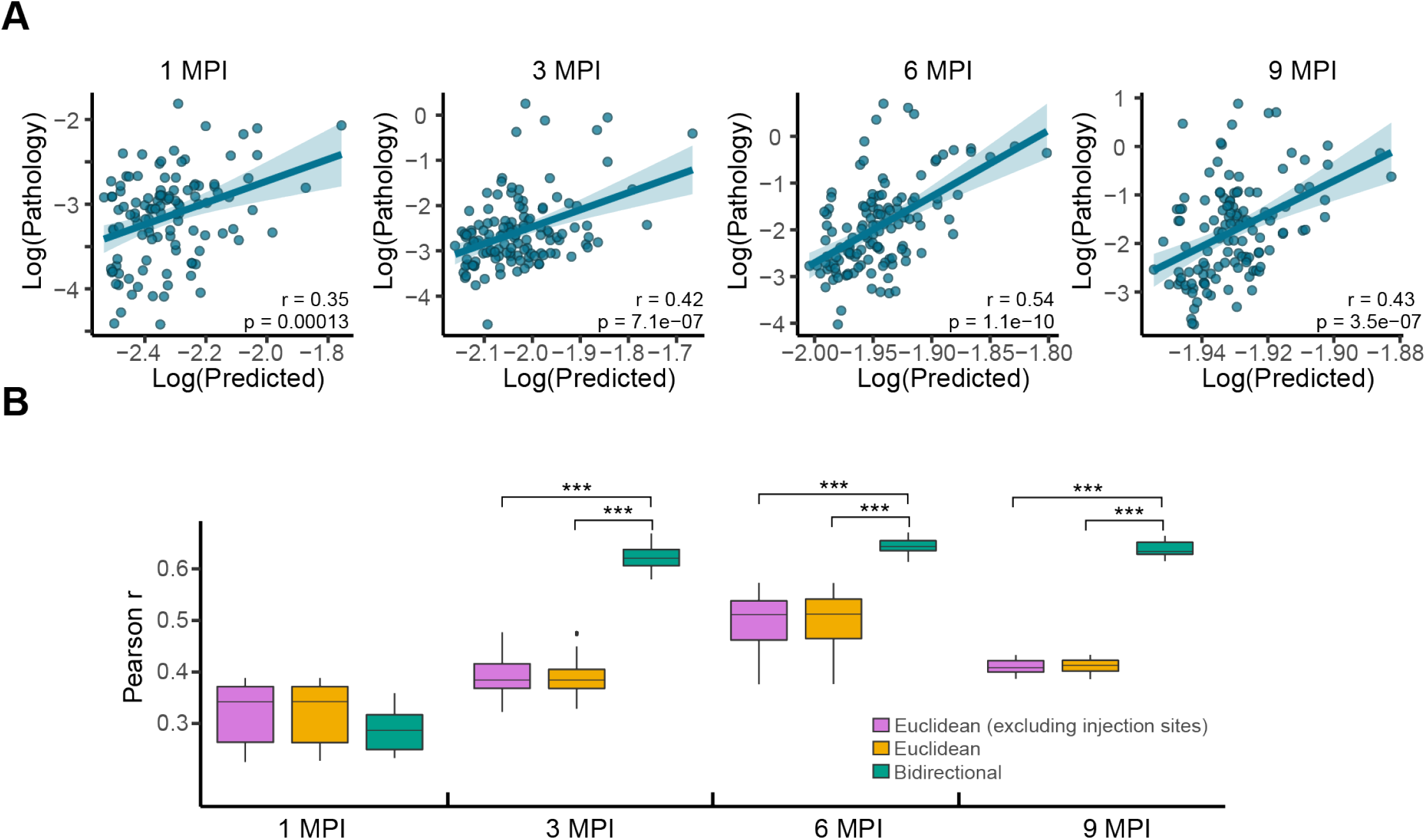
Effect of injection site outliers in Euclidean distance model on model fit, Related to Figure 4. (A) Predictions of log tau pathology (x-axis) from spread models based on Euclidean distance, plotted against log actual regional tau pathology values (y-axis) at 1, 3, 6, and 9 MPI. In these models, injection sites were removed from the sample prior to time constant optimization. The solid lines represent the lines of best fit, and the shaded ribbons represent the 95% prediction intervals. The r and p values for the Pearson correlation between model fitted values and observed pathology are noted on the plots. (B) Distributions of model fits in 100 held-out samples using a Euclidean distance model with injection sites exclude, a Euclidean distance model with all regions, and a bidirectional anatomical model with all regions. The performance of the Euclidean distance model does not depend on the inclusion or exclusion of injection sites, and the anatomical bidirectional model outperforms the Euclidean distance model in either case (***p<0.01).

**Figure S6.**
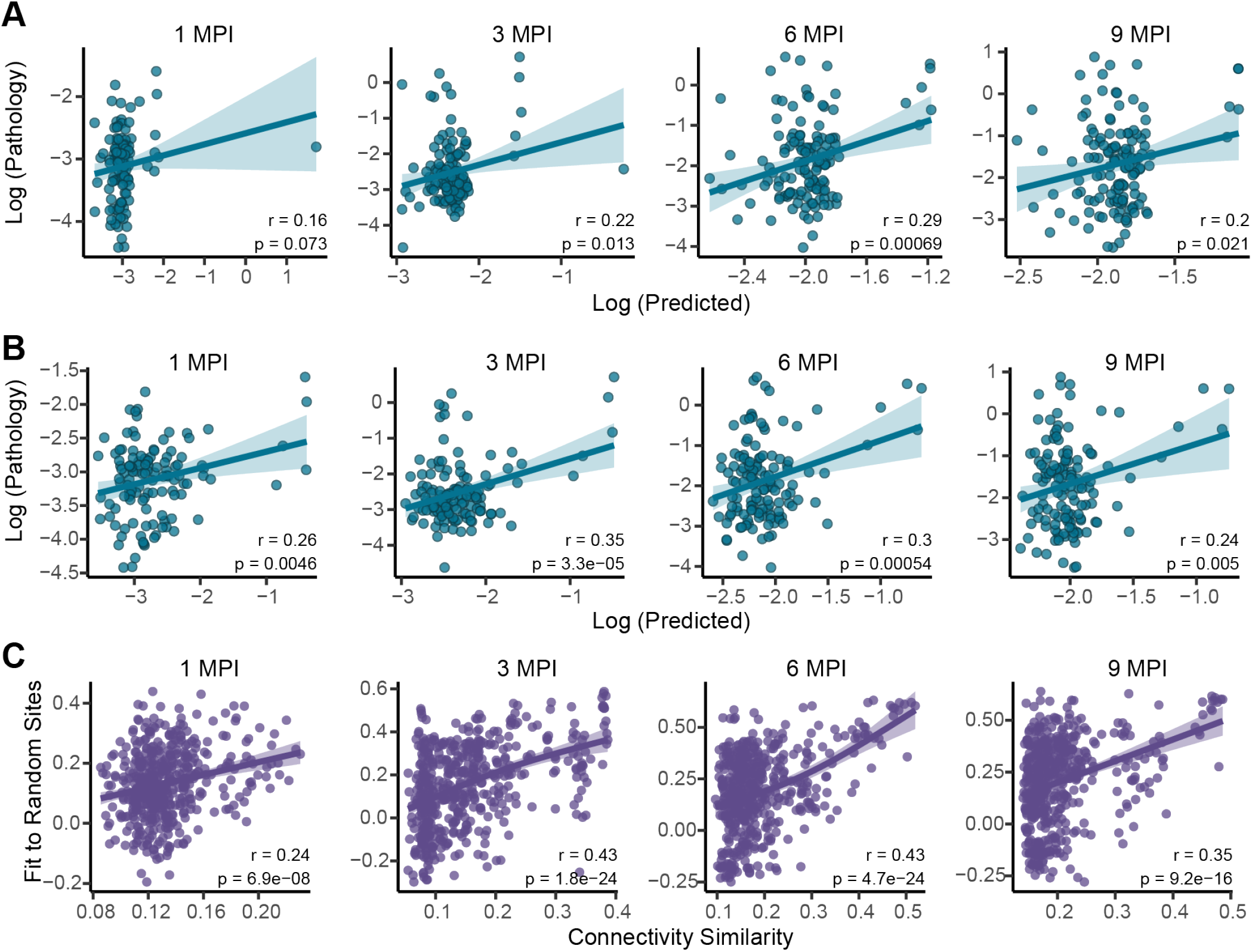
Tau pathology spread is predicted by diffusion through neuroanatomical connectivity, Related to Figure 4. Fits obtained using retrograde spread along a rewired network that preserves in-degree (B) or out-degree (C). (D) Alternate seed fit is partially explained by in projection similarity, out projection similarity, distance to injection site, and distance between alternate seed sites. The solid lines represent the lines of best fit, and the shaded ribbons represent the 95% prediction intervals. The r and p values for each linear regression are noted on the plots.

**Figure S7.**
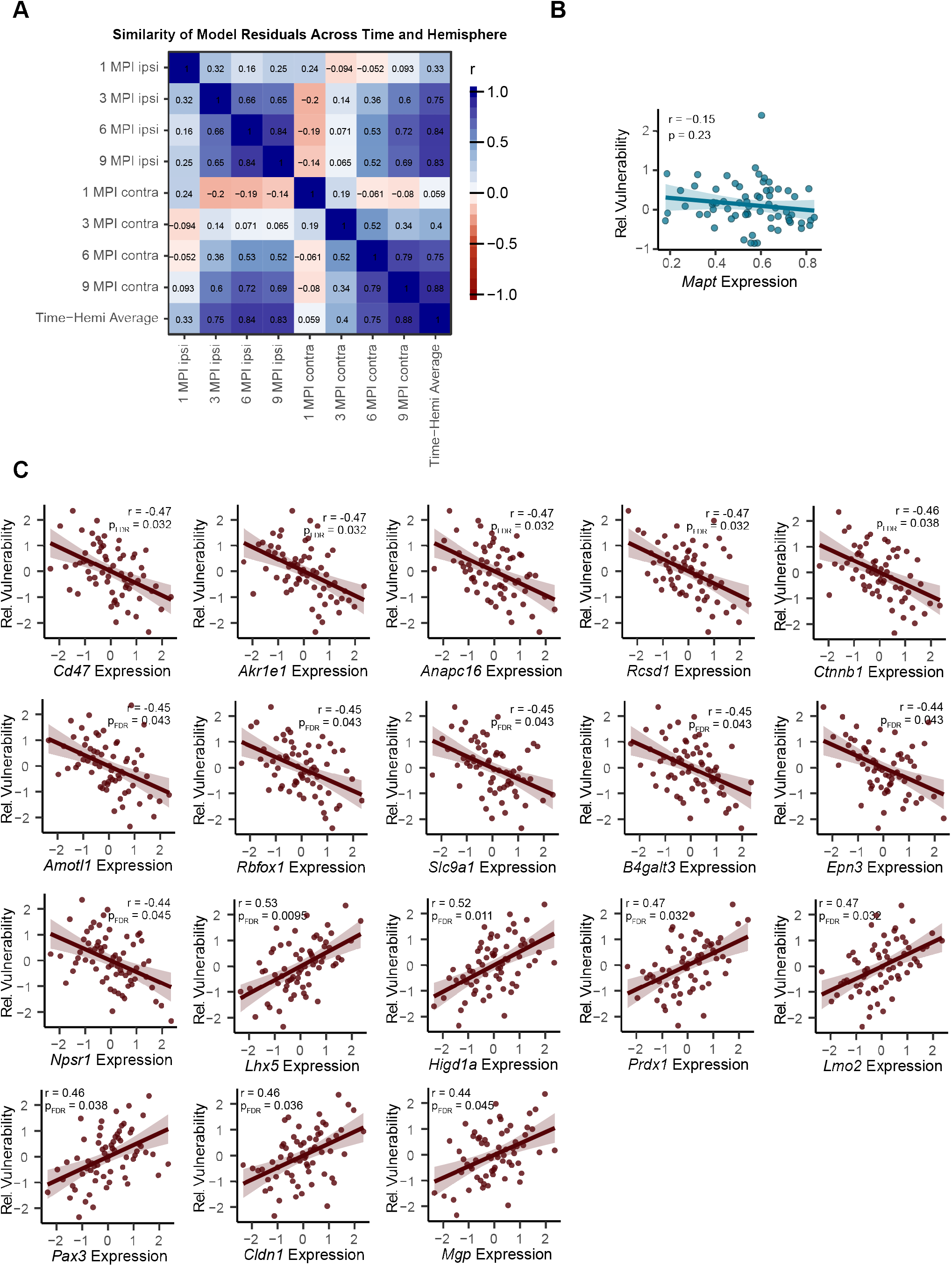
Assessment of model residuals as a proxy for regional vulnerability, Related to Figure 5. (A) Spatial similarity of model residuals between each hemisphere-time point combo. (B) Relative regional vulnerability is plotted as a function of *Mapt* expression. (C) Relative regional vulnerability is plotted as a function of gene expression patterns (FDR corrected p<0.05 cut off for inclusion). For panels (B) and (C), the solid line represents the line of best fit, and the shaded ribbons represent the 95% prediction intervals. The r and p values for the Pearson correlation between vulnerability and gene expression are noted on the plots.

**Figure S8.**
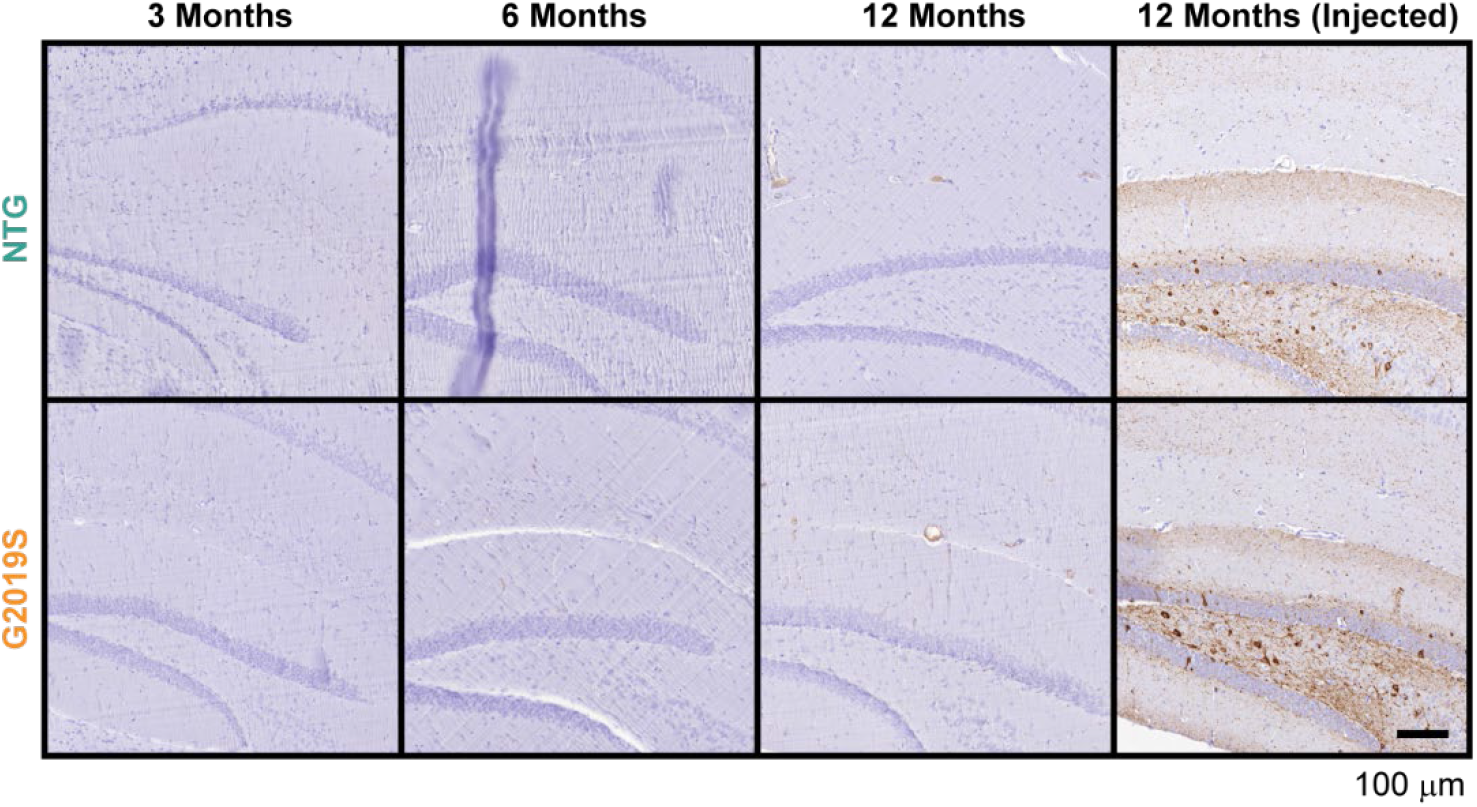
LRRK2^G2019S^ mice do not have tau pathology without pathogenic tau injection, Related to Figure 7. NTG and LRRK2^G2019S^ mice at 3, 6, or 12 months of age with no injection were assessed by immunohistochemistry for tau pathology in the hippocampus. In parallel, sections from NTG or LRRK2^G2019S^ mice injected with pathogenic tau were stained. These mice were 9 MPI or 12 months of age. No tau pathology was noted in non-injected mice. Scale bar = 100 μm.

**Figure S9.**
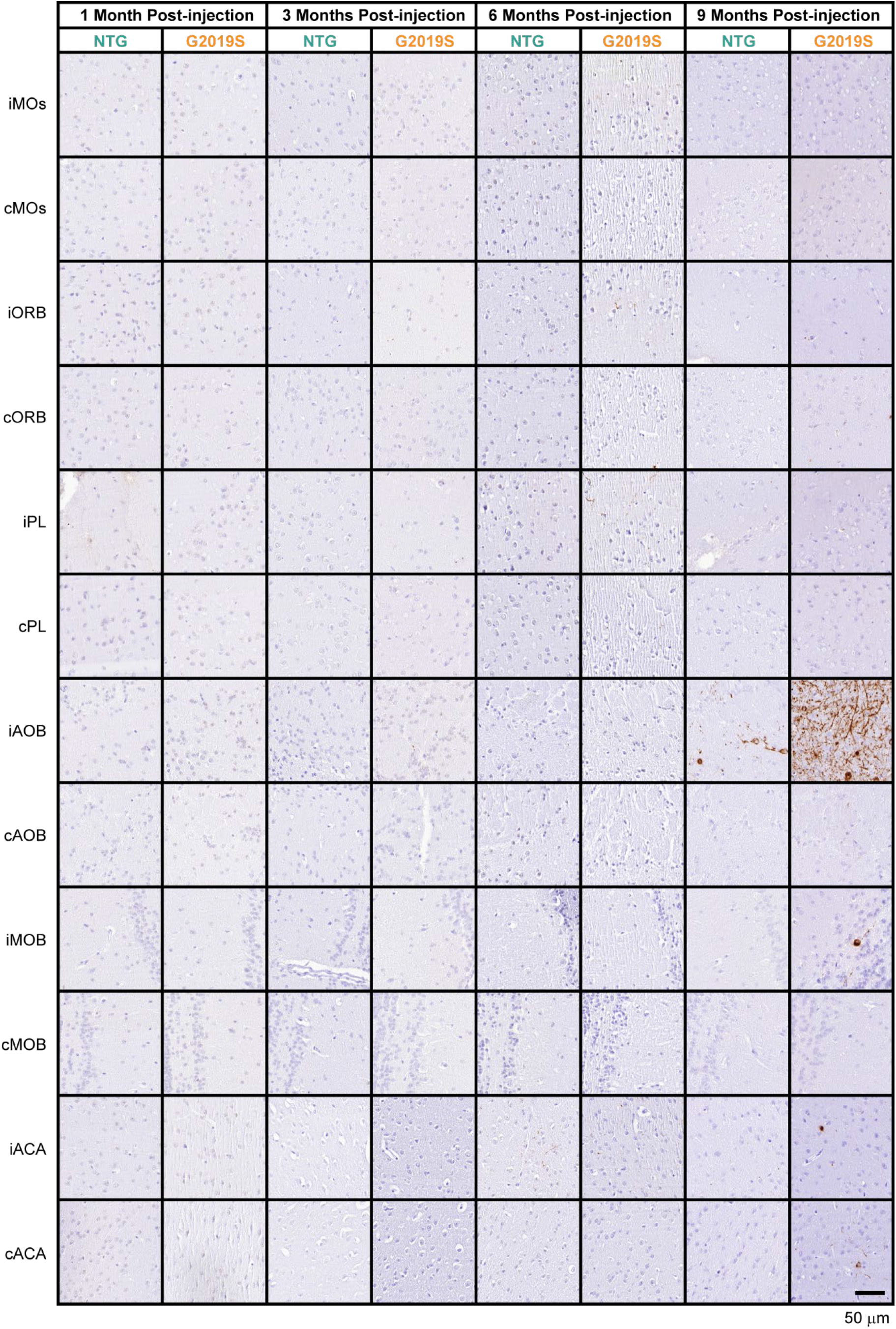
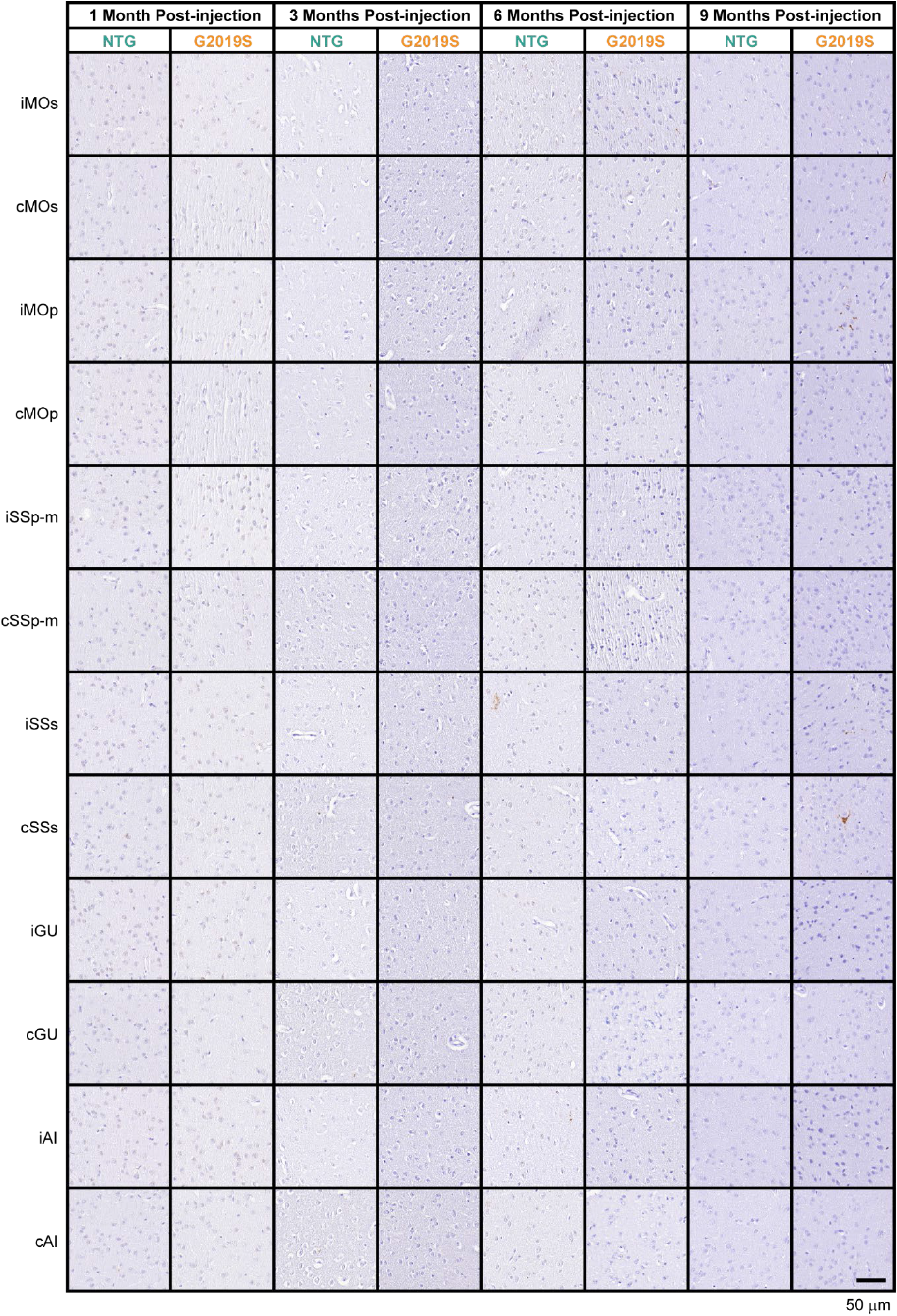
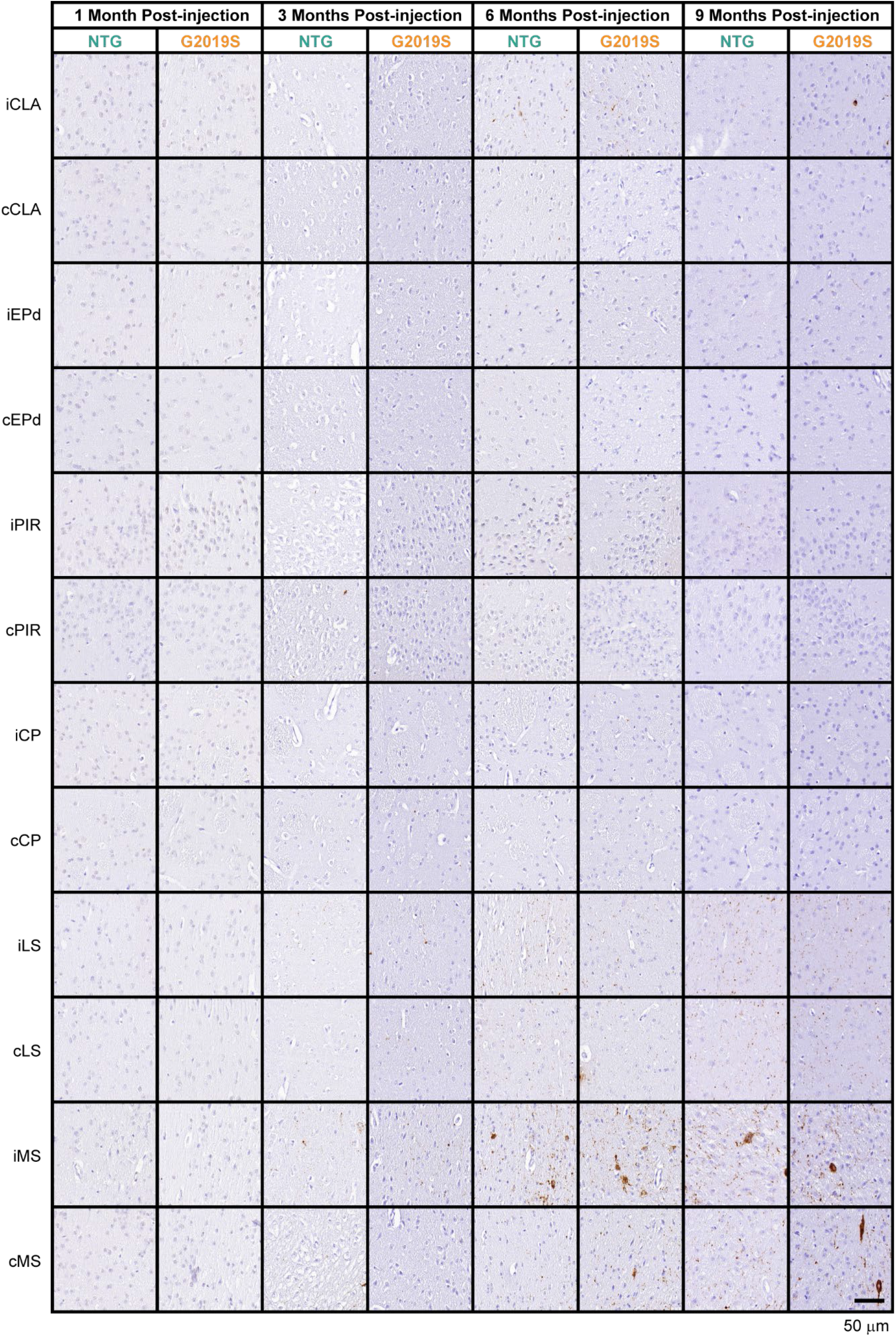
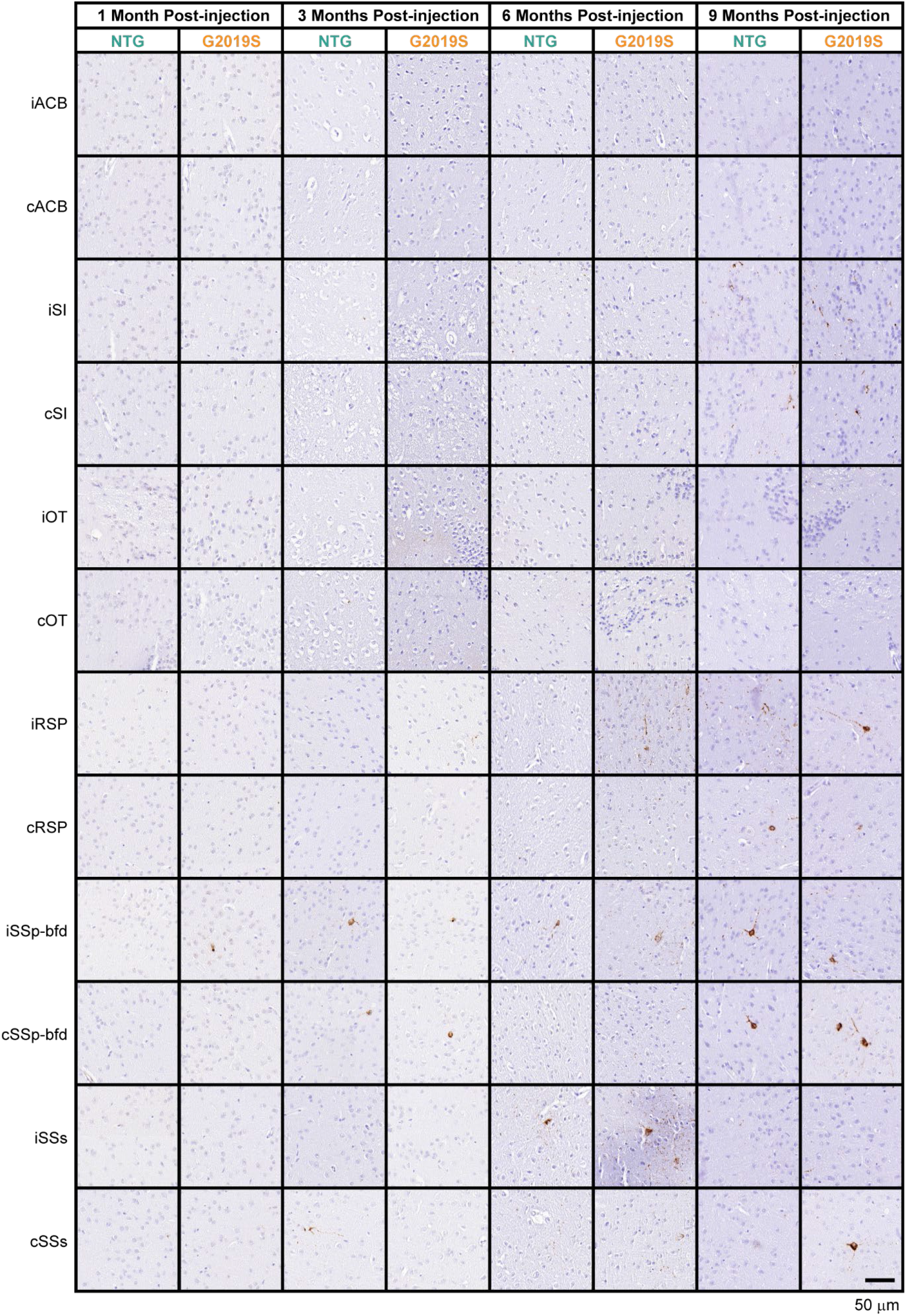
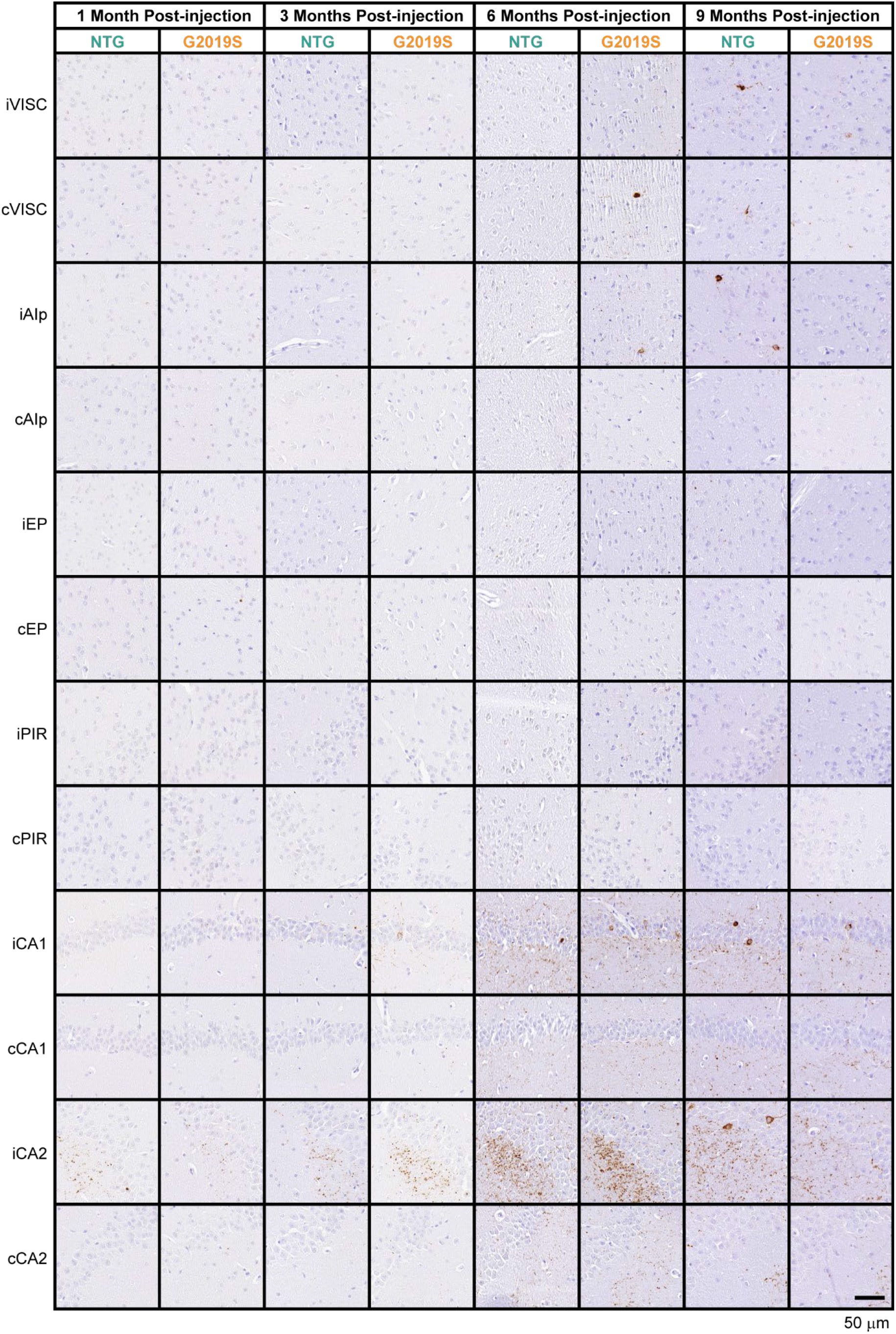
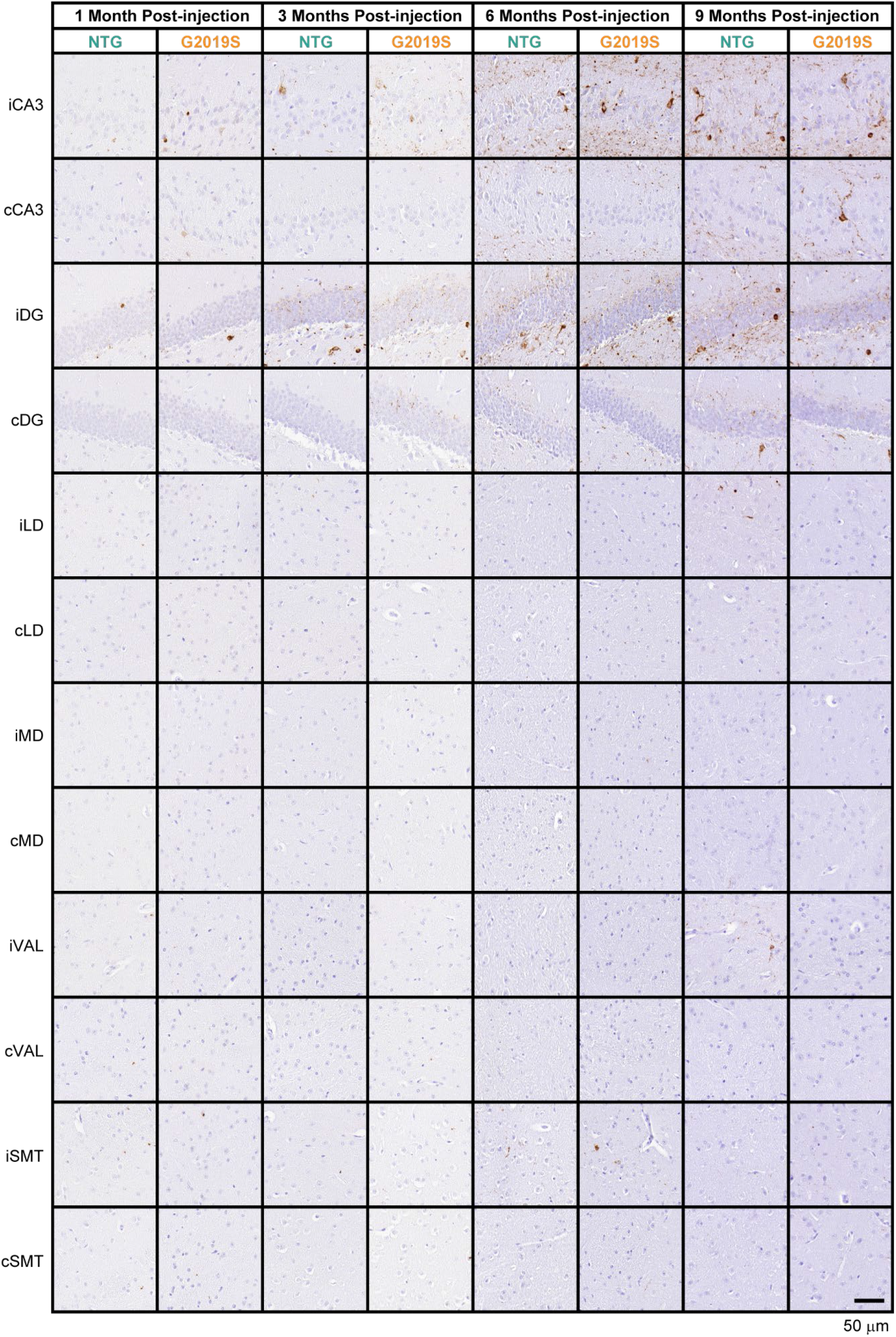
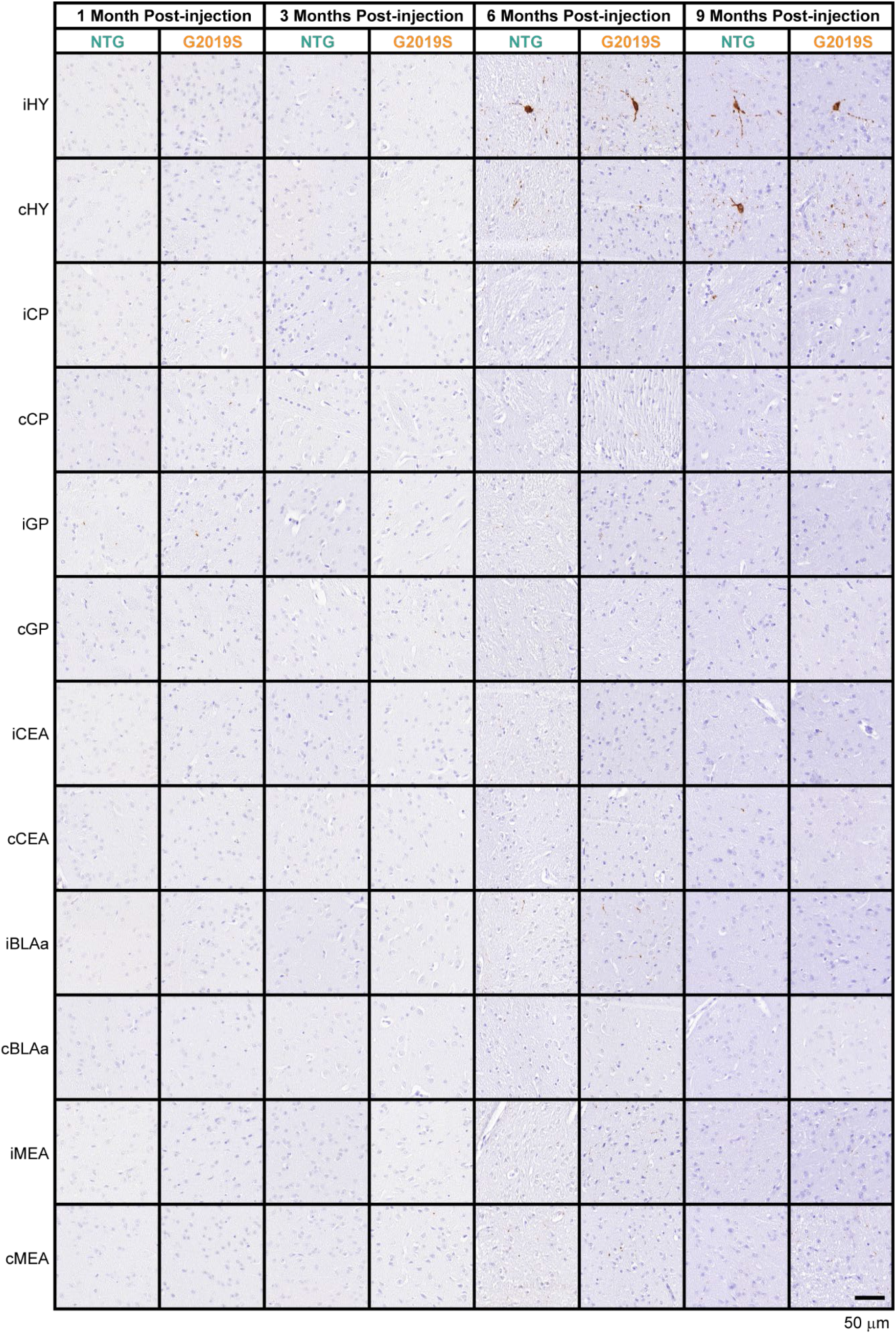
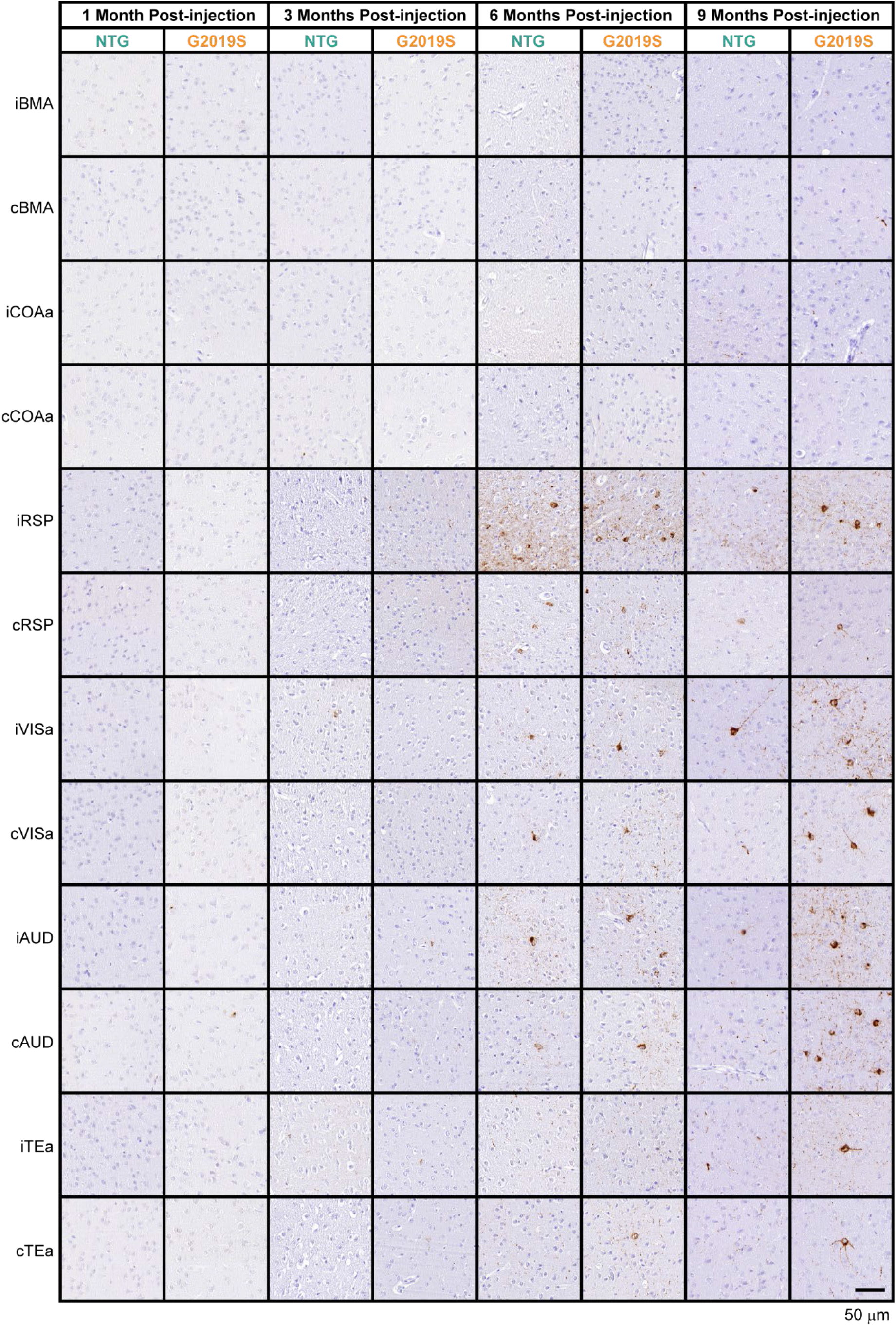
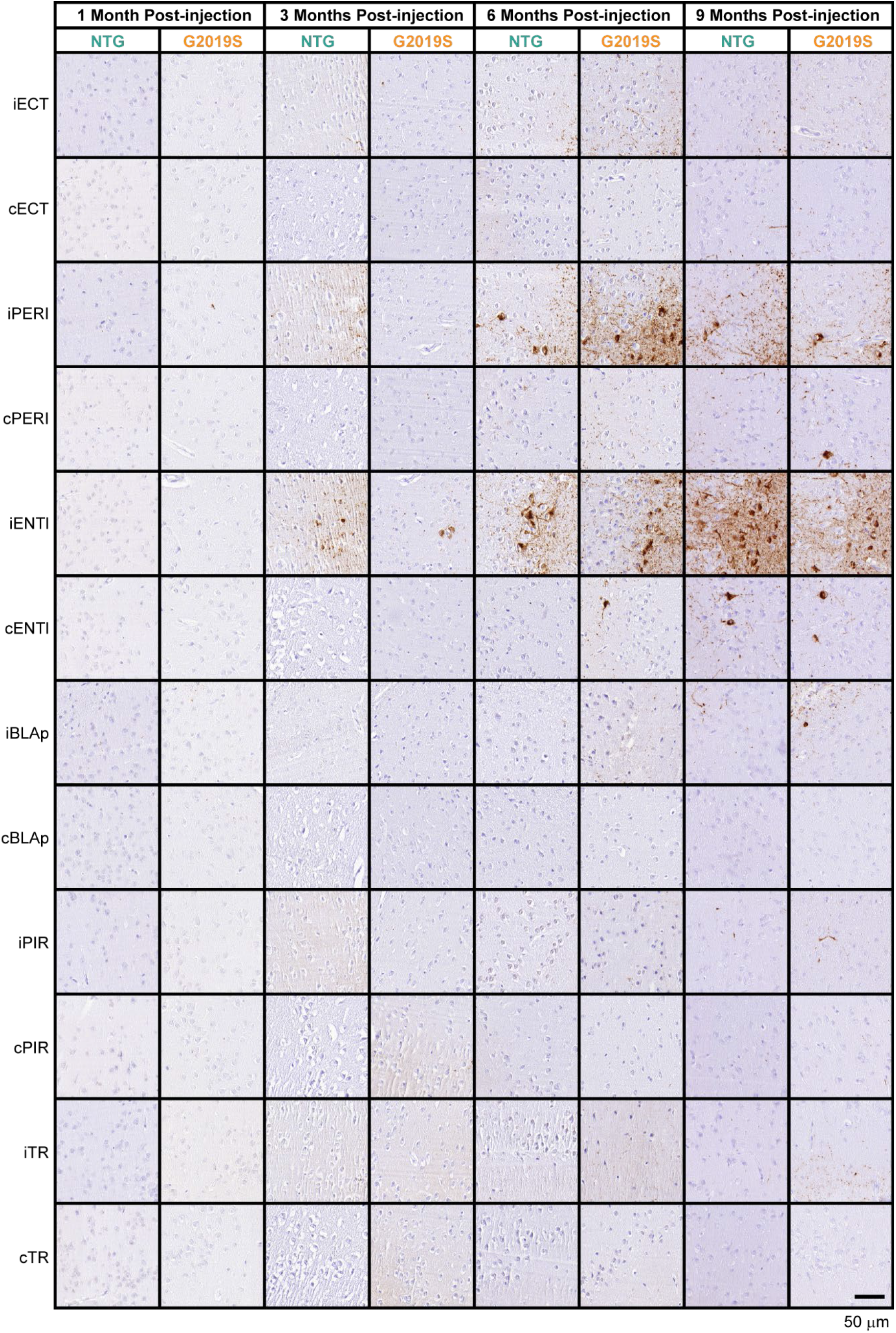
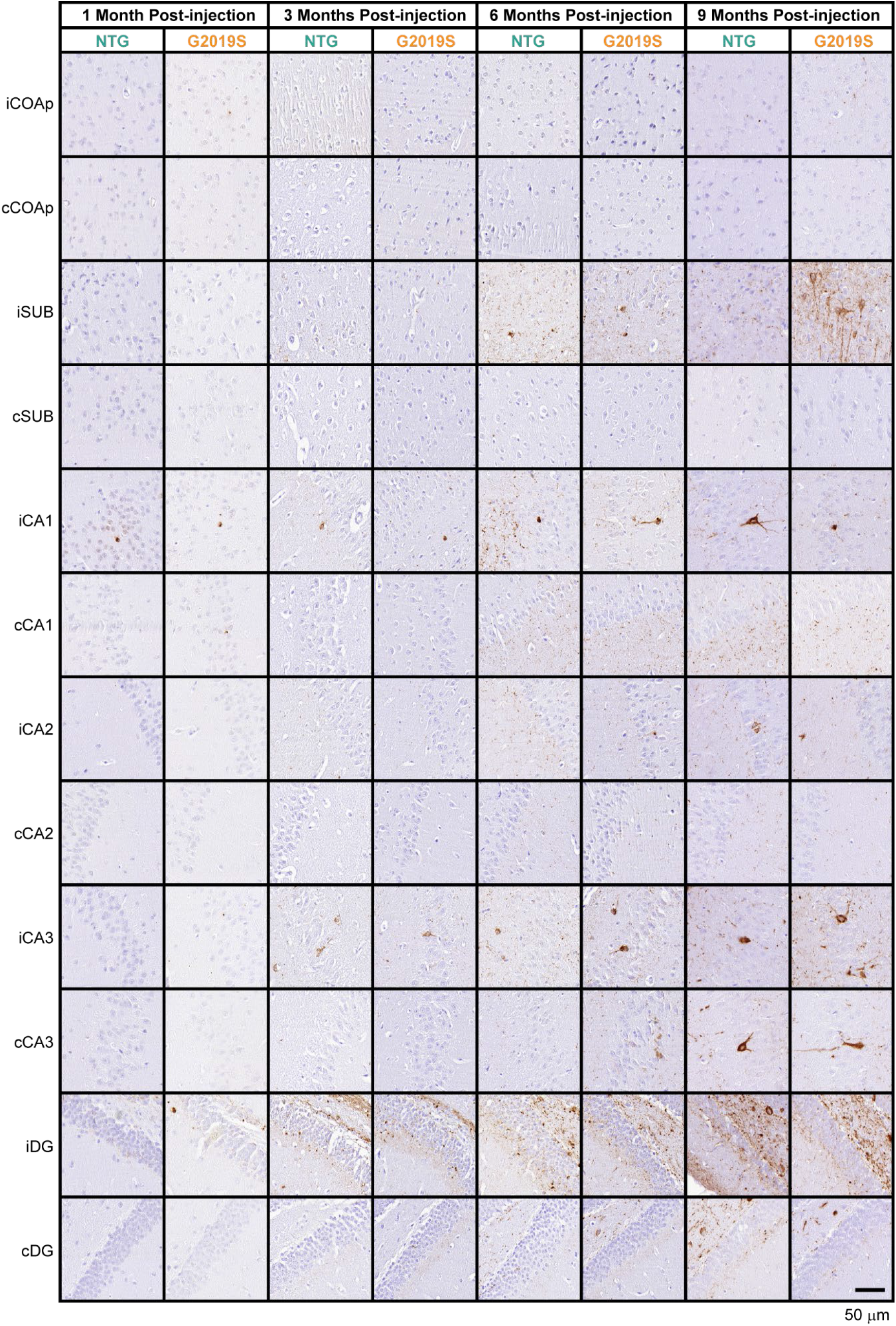
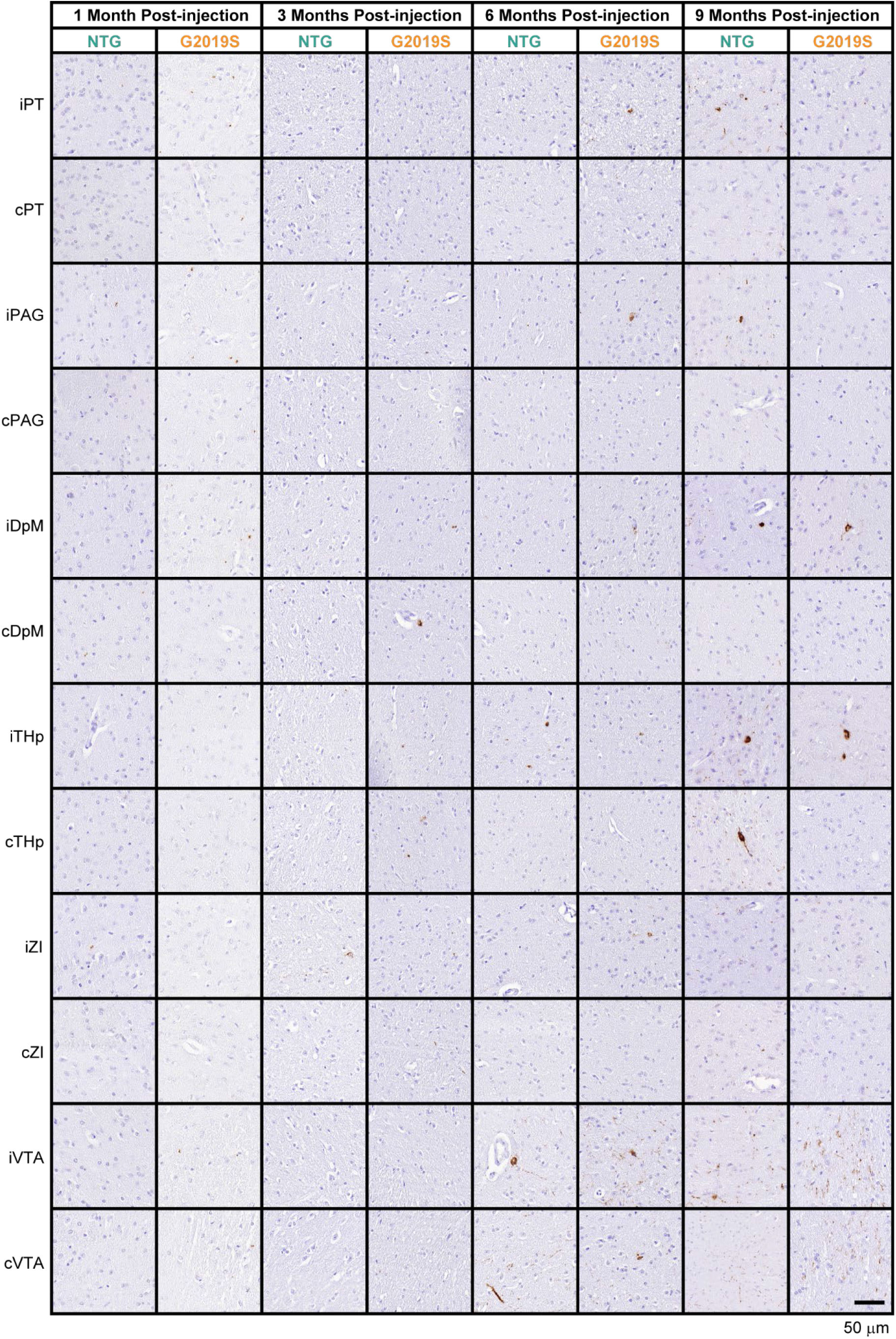
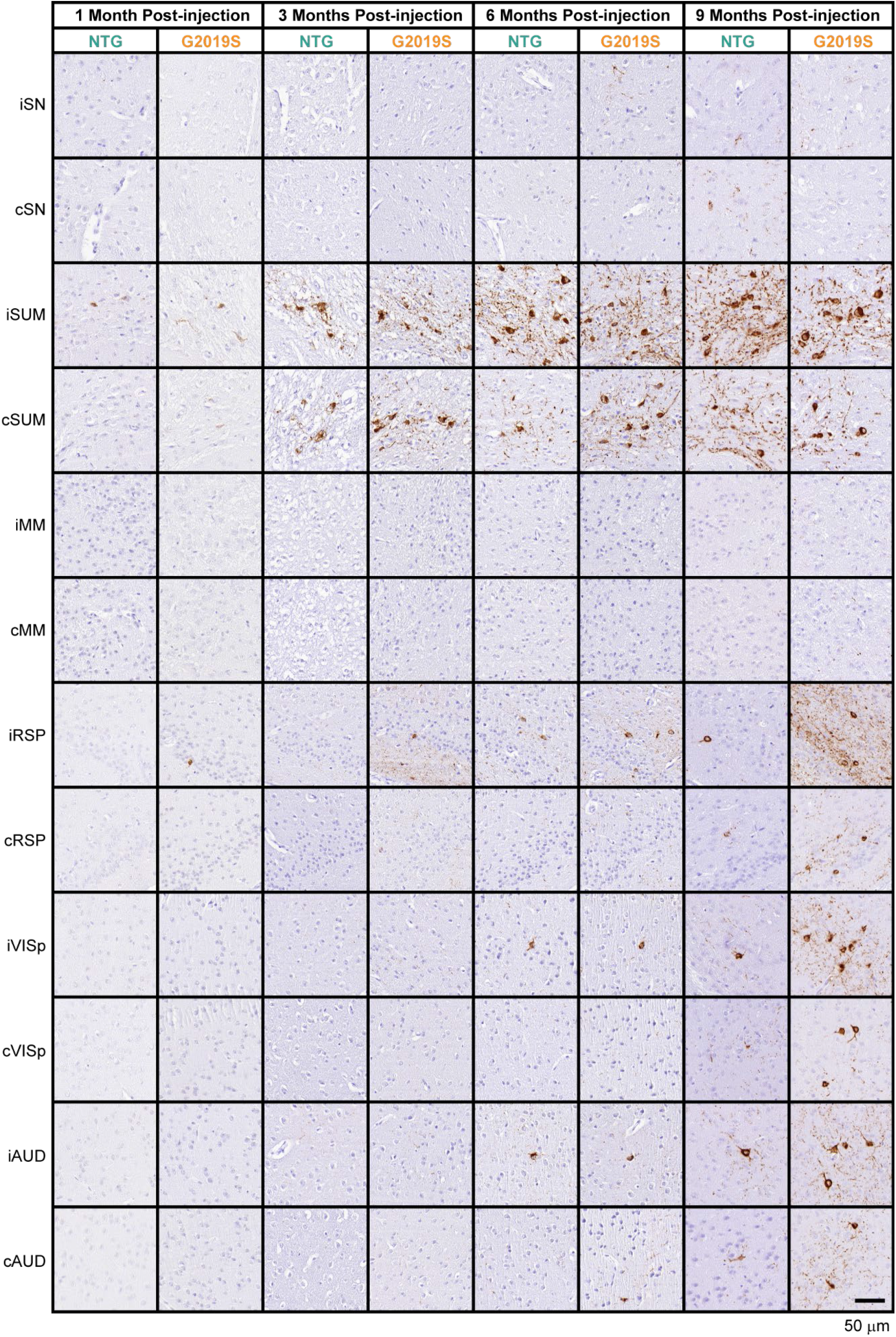
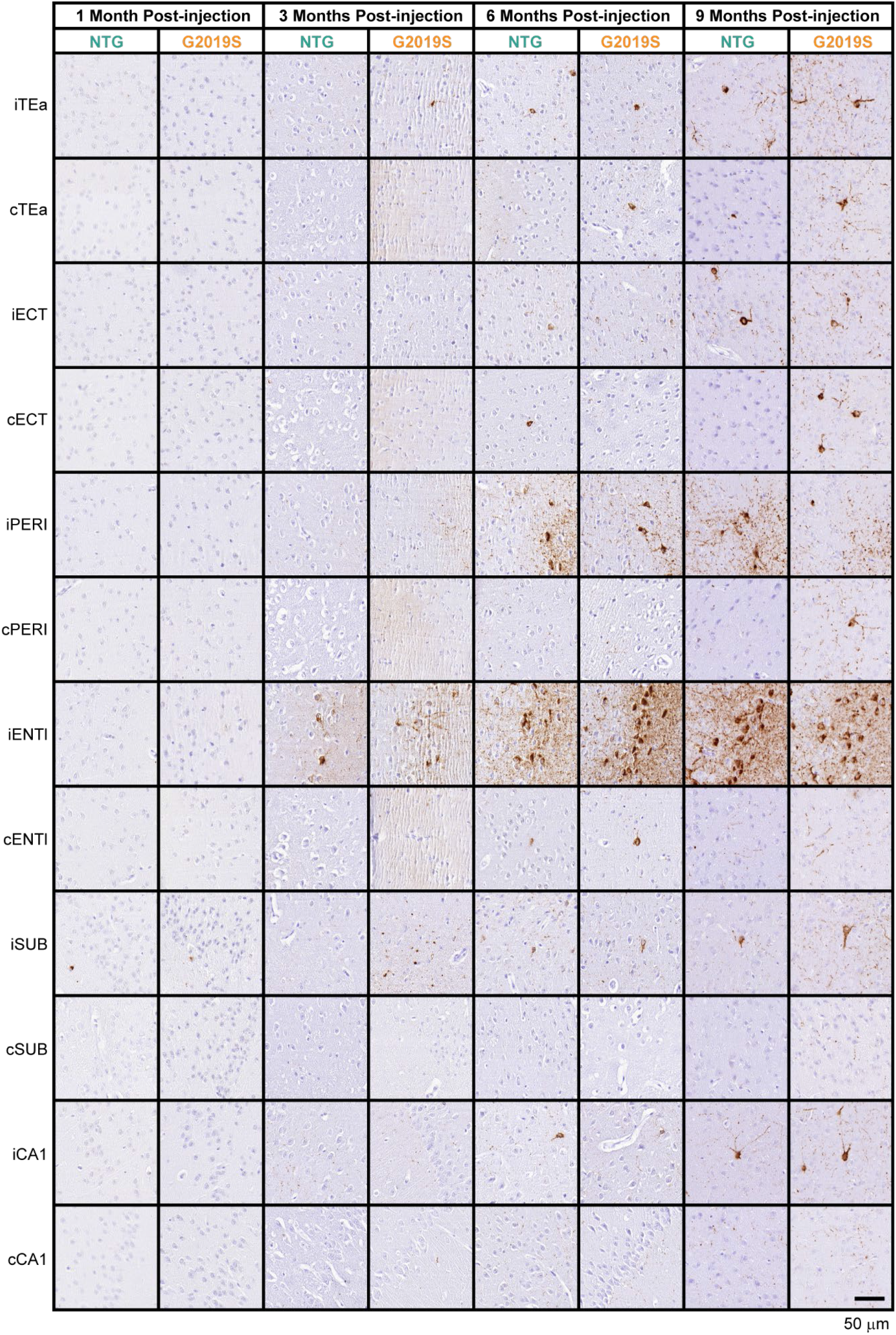
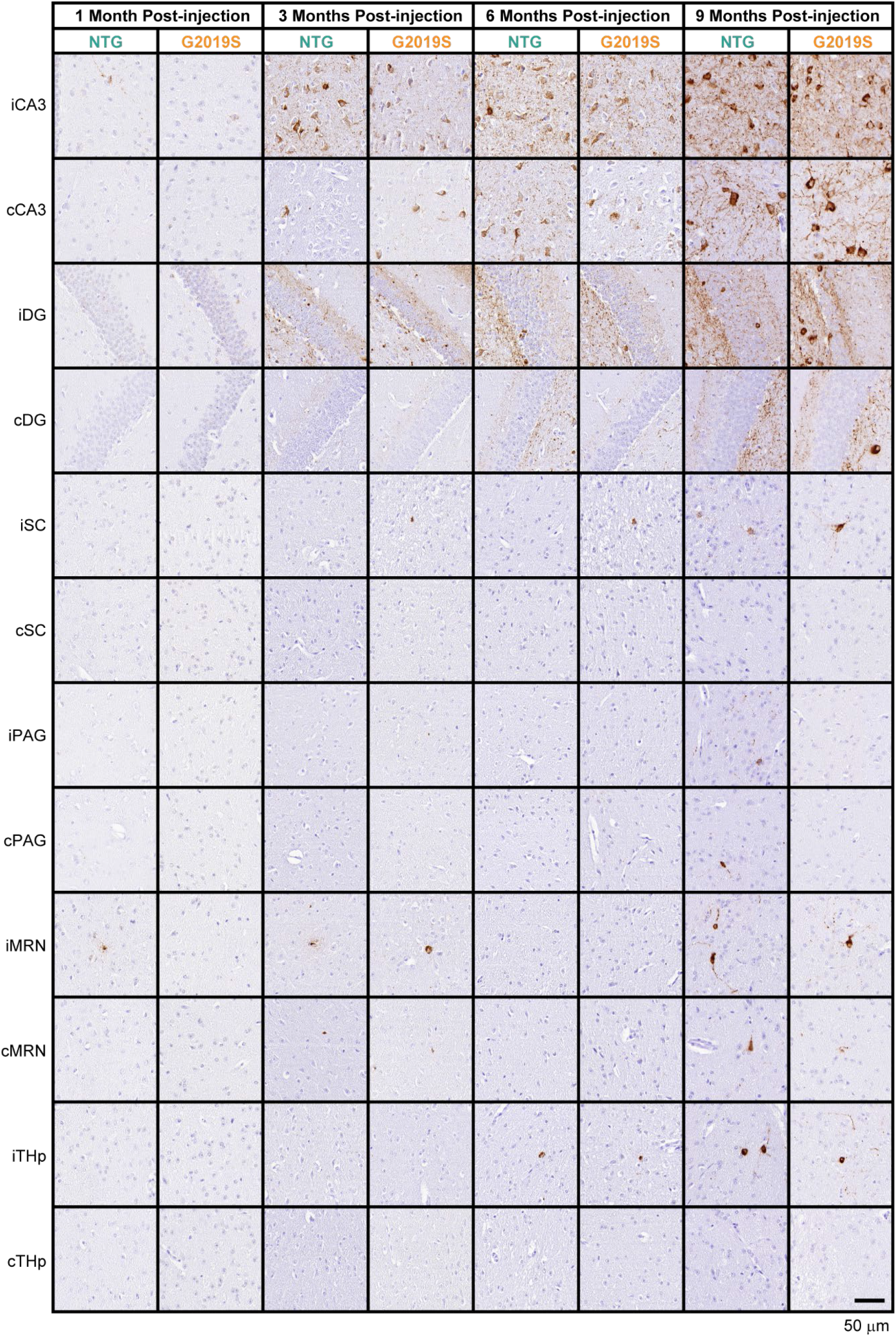
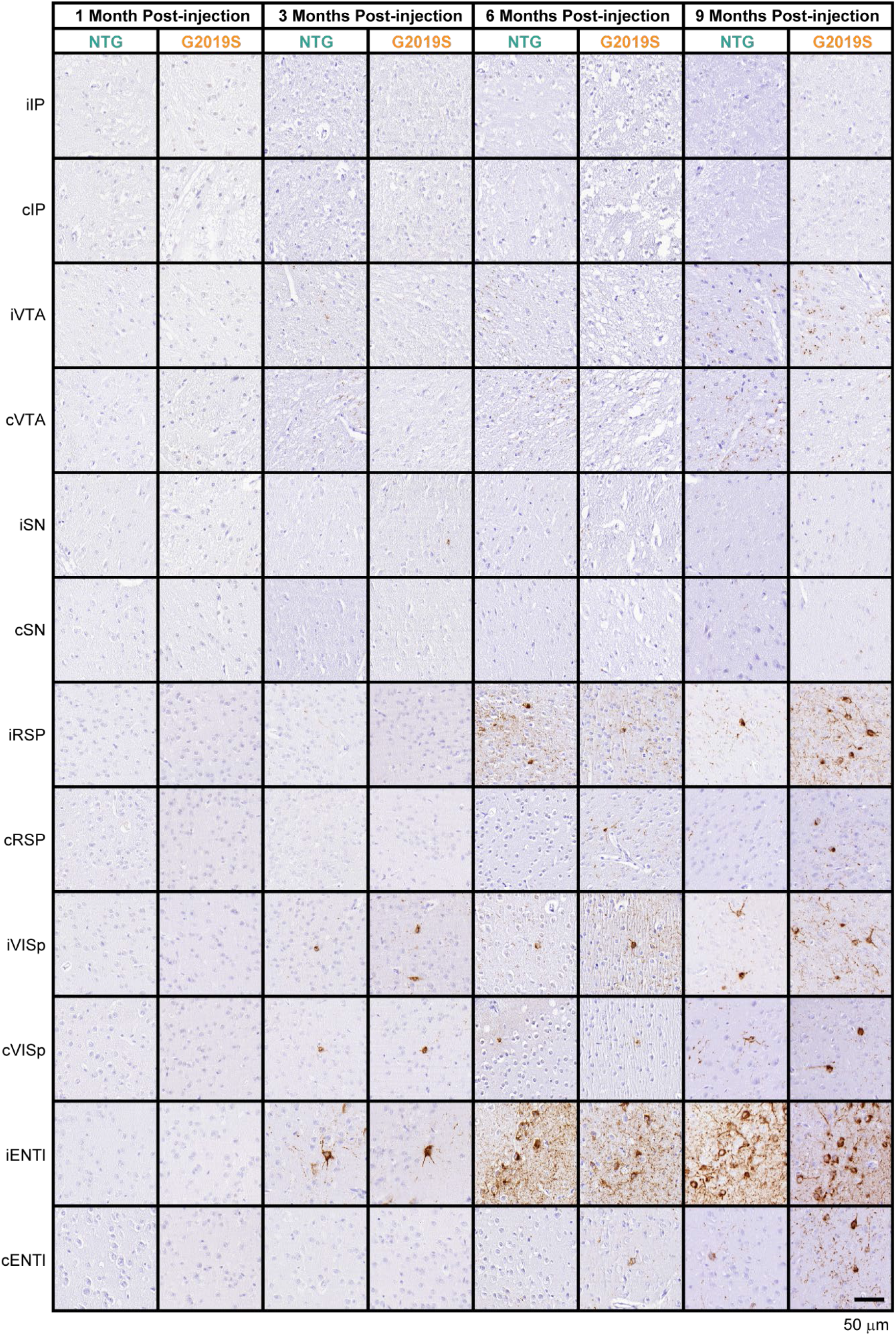
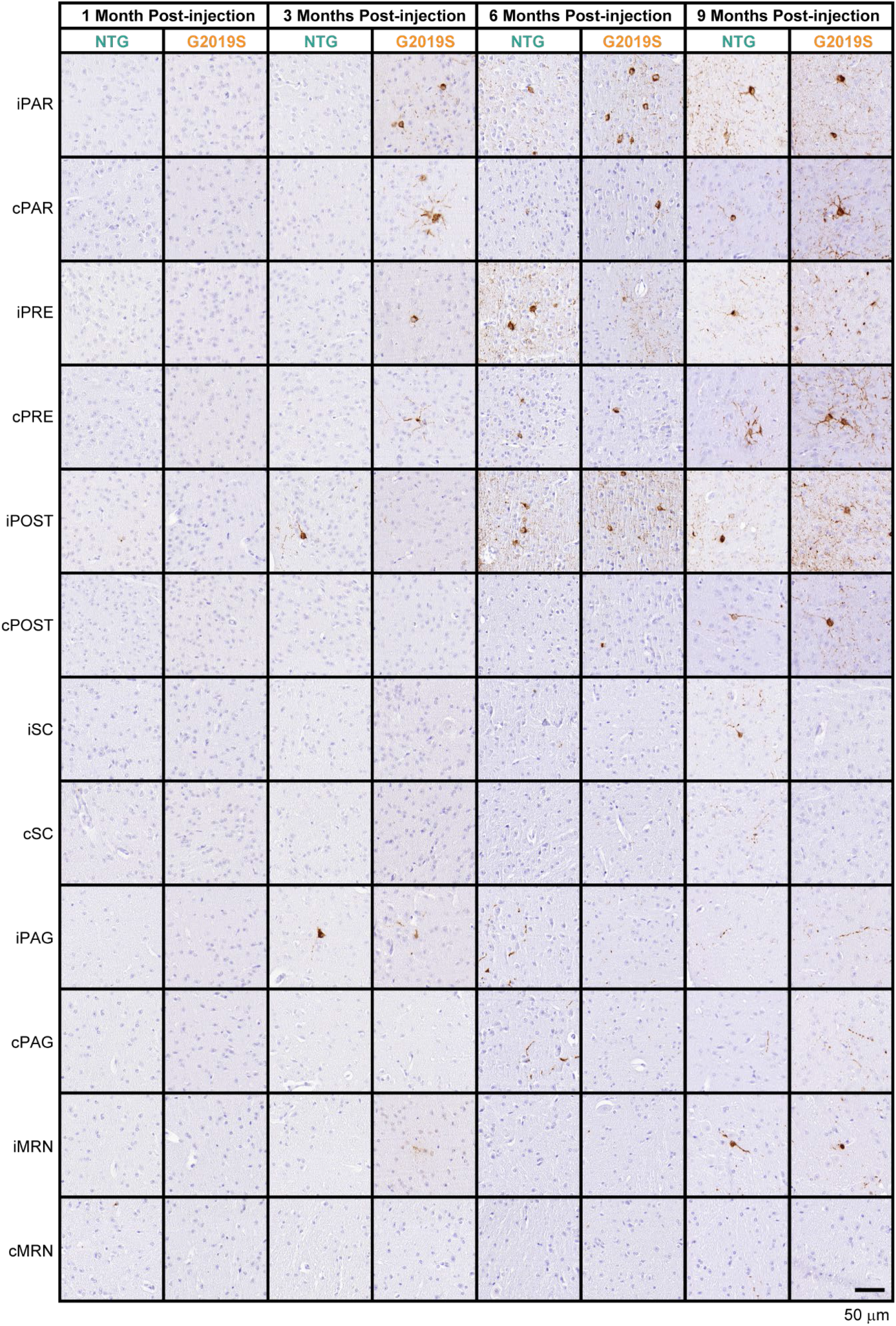
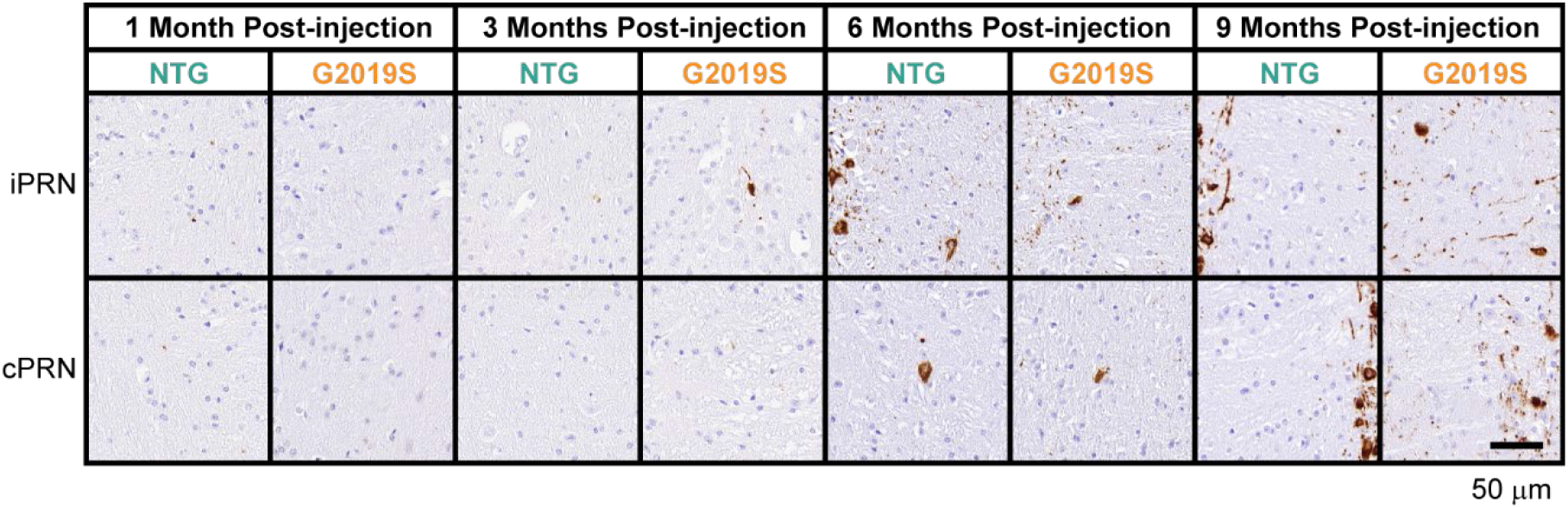
Representative pathology images from all quantified regions. pS202/T205 tau (AT8) staining (scale bars = 50 μm). See Supplemental Fig. 1 for region designations (“I” precedes ipsilateral regions and “c” precedes contralateral regions). Some regions are represented multiple times if they were quantified in multiple coronal sections.

